# Accurate models of substrate preferences of post-translational modification enzymes from a combination of mRNA display and deep learning

**DOI:** 10.1101/2022.02.14.480467

**Authors:** Alexander A. Vinogradov, Jun Shi Chang, Hiroyasu Onaka, Yuki Goto, Hiroaki Suga

## Abstract

Promiscuous post-translational modification (PTM) enzymes often display non-obvious substrate preferences by acting on diverse yet well-defined sets of peptides and/or proteins. Thorough understanding of substrate fitness landscapes for promiscuous PTM enzymes is important because they play key roles in many areas of contemporary science, including natural product biosynthesis, molecular biology and biotechnology. Here, we report the development of an integrated platform for accurate profiling of substrate preferences for PTM enzymes. The platform features a combination of i) mRNA display with next generation sequencing as an ultrahigh throughput technique for data acquisition and ii) deep learning for data analysis. The high accuracy (>0.99 in each of two studies) and generalizability of the resulting deep learning models enables comprehensive analysis of enzymatic substrate preferences. The models can be utilized to quantify fitness across sequence space, map modification sites, and identify important amino acids in the substrate. To benchmark the platform, we perform substrate specificity profiling of a Ser dehydratase (LazBF) and a Cys/Ser cyclodehydratase (LazDEF), two enzymes from the lactazole biosynthesis pathway. In both studies, our results point to highly complex enzymatic preferences, which, particularly for LazBF, cannot be reduced to a set of simple rules. The ability of the constructed models to dissect and analyze such complexity suggests that the developed platform can facilitate the wider study of PTM enzymes.

## Introduction

Enzymes which perform post-translational modification (**PTM**) of peptides and proteins often display non-trivial preferences by acting on a wide range of substrates. The nuanced and, in many cases, poorly understood nature of substrate recognition and engagement by PTM enzymes has come under scrutiny in several contexts. For example, during the biosynthesis of ribosomally synthesized and post-translationally modified peptides (**RiPP**s),^1,2^ notably lanthipeptides^3,4^ and cyanobactins,^5^ a single set of PTM enzymes can modify disparate substrates to assemble multiple natural products.^6,7^ Catalytic promiscuity of RiPP biosynthetic enzymes suggests numerous bioengineering applications,^8–11^ and accordingly, much effort has been dedicated to understanding the molecular basis for such behavior.^12–17^ Likewise, in human biology, dense PTM networks controlled by hundreds of promiscuous enzymes orchestrate virtually every aspect of cell behavior, and thus, investigating how PTM enzymes discriminate their substrates is integral to forming a holistic appreciation of cellular processes.^18–20^

Substrate specificity profiling studies are a natural first step when studying catalysis by promiscuous PTM enzymes. Numerous approaches developed over the years^21–25^ enable streamlined analysis of substrate fitness landscapes, but every method comes with its own limitations. Platforms based on the screening of synthetic peptide microarrays^26–30^ and saturation mutagenesis approaches^31–33^ have relatively low throughput, and can suffer from limited generalizability and accuracy. For example, microarray-derived substrate preferences of sirtuin lysine deacetylases mismatch known cellular substrates.^34^ In vivo library construction methods, particularly yeast and phage display, offer a much higher throughput (up to ∼10^9^ peptides for testing compared to 10^3^-10^4^ for microarrays), but developing experimental schemes for phage/yeast display can be technically difficult, and these approaches to date have mainly focused on studying proteases.^35–37^

Inference from large amounts of data is another challenge common for high throughput methods. The de facto standard approach is the computation of position-wise amino acid enrichment scores (usually displayed as WebLogo sequence alignment plots),^38^ which over-compresses available information and inevitably loses the nuance. Machine learning/deep learning methods represent a promising alternative to the purely statistical treatment of data. Deep learning has in the recent years proven its ability to make meaningful generalizations in a variety of complex tasks, but it requires large amounts of clean, curated data to train and evaluate the models.^39,40^ To date, the substrate profiling studies which utilized deep learning were either data-limited, due to their reliance on peptide microarrays for data acquisition,^41^ or used heterogeneous datasets,^42–45^ which have led to models with modest predictive power.

Messenger RNA (mRNA) display-based enzyme profiling strategies^46,47^ have recently gained traction as a viable alternative to the established methods. As a fully in vitro platform, mRNA display can access combinatorial libraries of vast diversity (>10^12^ unique peptides).^48,49^ The technique also allows for elaborate manipulation of the libraries (extensive genetic code reprogramming, affinity purification, chemical labelling, etc.) and therefore, supports the development of workflows inaccessible to in vivo methods. Still, mRNA display approaches have thus far revolved around single-point saturation mutagenesis,^46,47^ and as such, have typically profiled only hundreds of enzyme substrates at once, not taking full advantage of the available library diversity.

Here, we report the development of a general platform for assaying substrate fitness landscapes of PTM enzymes (Fig. 1). Our approach integrates mRNA display as the data-generating engine with deep learning workflows to analyze and learn from the resulting data. Using two RiPP biosynthetic enzymes catalyzing distinct reactions, we demonstrate that mRNA display-based substrate selections can provide large amounts of clean, labelled data to train supervised deep learning classifiers of enzymatic substrate preferences. The resulting models accurately predict substrate fitness from primary sequence and generalize well across the peptide sequence space. The models have a degree of interpretability that allows for mapping of modification sites and identification of important residues in the substrate. Altogether, we believe that the described pipeline is a powerful tool for studying the dynamics of PTM enzyme/substrate interactions.

**Figure 1.**
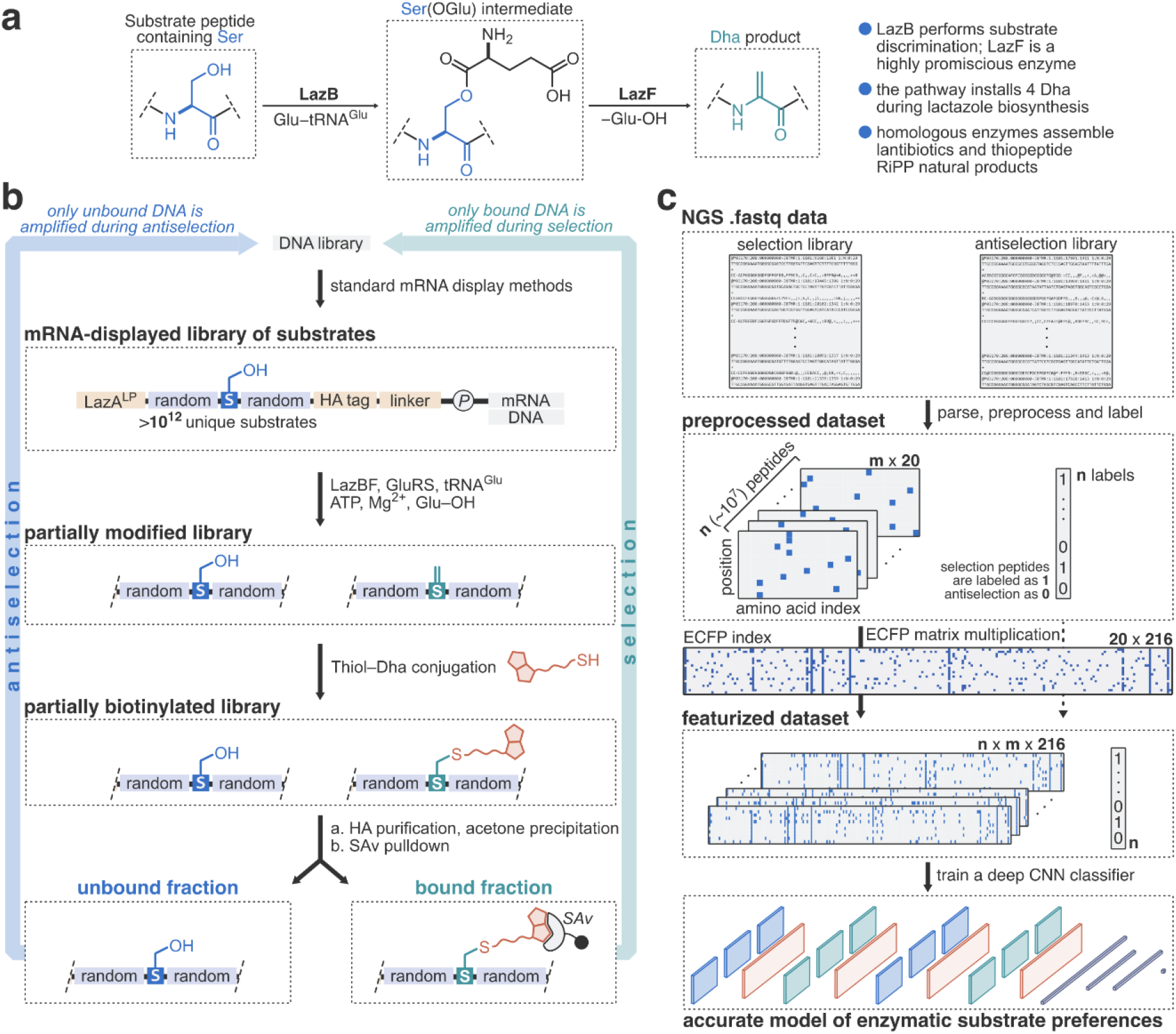
An overview of the workflow for the profiling of LazBF substrate preferences. **a**) Chemical reaction catalyzed by LazBF. **b**) Schematic overview of mRNA display-based selection/antiselection setups. For the full protocol, see S.I. 2.3. Ⓟ refers to the puromycin linker used to display the peptides onto cognate mRNAs. Both selection and antiselection assays can be repeated several times to produce libraries of progressively increasing (or decreasing) substrate fitness. **c**) Schematic overview of the data analysis pipeline. NGS selection and antiselection datasets are parsed, preprocessed and labelled. Peptides are represented as positionally encoded matrices of EFCPs, and a supervised CNN classifier is trained on the resulting data to produce models of LazBF substrate preferences. For the complete description of the data analysis pipeline, see S.I. 2.5.

## Results and discussion

### Development of the mRNA display scheme for LazBF profiling

For this study, we focused on PTM enzymes participating in the biosynthesis of lactazole A,^50^ a natural product belonging to the thiopeptide family of RiPPs.^51^ The compound is a promising bioengineering target because its five biosynthetic enzymes (LazBCDEF) can convert a wide variety of sequence-randomized precursor peptides to lactazole-like thiopeptides.^47,52^ LazBF, a split Ser dehydratase homologous to class I lanthipeptide synthetases (Fig. 1a),^3^ plays a central role during lactazole biosynthesis because its operation to install four dehydroalanine (**Dha**) residues into precursor peptide LazA requires precise timing and selectivity.^53^ Mechanistically, LazBF operates via a two-step process akin to class I Ser/Thr dehydratases.^54,55^ The glutamylation domain (LazB) binds the N-terminal leader peptide (**LP**) region of LazA (**LazA**^**LP**^) and promotes Ser glutamylation in the downstream core peptide (**CP**) section using Glu-tRNA^Glu^ as the acyl donor. In the second step, the elimination domain (LazF) catalyzes a retro-Michael reaction in the Ser(OGlu) intermediate to yield the Dha-containing product.^56^ Although preliminary enzyme characterization indicated that LazBF prefers a Trp residue in position +1 relative to the modification site, the enzyme also displayed more elaborate preferences which eluded generalization.^47,53^ Here, we have sought to develop an mRNA display/deep learning-based platform for comprehensive profiling of LazBF substrate fitness landscapes.

We envisioned training a supervised learning classifier that could predict the fitness of LazBF substrates from their primary sequence. To that end, the acquisition of two mRNA display datasets (one corresponding to substrates of high fitness and another for peptides which are not dehydrated by LazBF) was necessary. We anticipated that the treatment of a diverse library of mRNA-displayed peptides with LazBF would dehydrate some, but not all, library members (Fig. 1b). The modified peptides, i.e., those containing a Dha residue, are reactive toward thiols^57^ and can be selectively conjugated to a biotinylated probe (biotin-PEG_2_-SH; Fig. S1d). The labelling reaction enables the separation of modified and unmodified substrates using streptavidin (**SAv**) pulldown, which selectively isolates biotinylated products. The subsequent PCR amplification of either the SAv-bound or unbound fraction recovers DNA libraries encoding peptides of increased or decreased fitness, respectively. Iterative repetition of this process should produce increasingly enriched peptide populations. During a “selection”, SAv-bound DNA is amplified to enrich for substrates of high fitness, while an “antiselection” recovers the unbound DNA fraction to generate a dataset of poor substrates.

To establish the assay, we designed an mRNA library encoding peptides bearing the LazA^LP^ sequence followed by a randomized CP, HA tag for affinity purification and a flexible C-terminal linker (library 5S5; Fig. 2). Every CP contained a potentially modifiable Ser residue flanked by five random amino acids on either side (theoretical diversity: 1·10^13^ sequences) to establish substrate recognition requirements around the dehydration site. Our preliminary experiments indicated that library 5S5 contained substrates of differential fitness. First, we selected three such peptides (bAP1–3, in order of decreasing fitness; Fig. S1) to establish the experimental conditions. The treatment of the peptides expressed by the flexible in vitro translation (**FIT**) system^58^ with 2 µM LazBF, 20 µM tRNA^Glu^ and 1 µM GluRS for 2 h led to the quantitative dehydration of bAP1, partial modification of bAP2, and virtually no reaction for bAP3 (Fig. S1a-c). Further incubation of the reaction products with 5 mM biotin-PEG_2_-SH at pH 8.5 on ice for 17 h resulted in selective and nearly quantitative biotinylation of Dha-containing peptides, indicating the feasibility of the envisioned experimental scheme (Fig. S1e).

**Figure 2.**
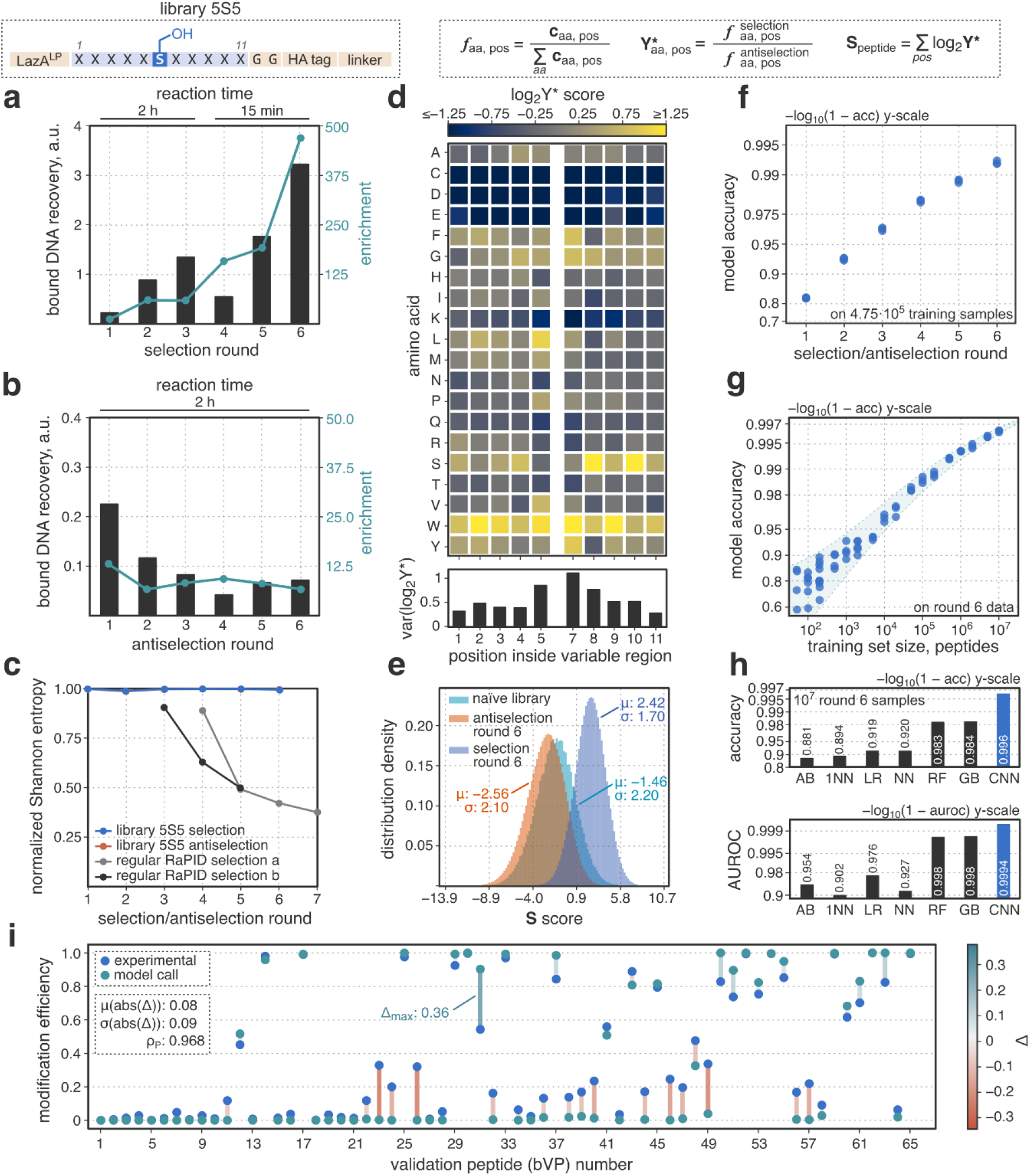
mRNA display profiling of LazBF leads to enriched peptide populations suitable for deep learning applications. **a, b**) Summary of the selection (a) and antiselection (b) experiments. Plotted are respective DNA recovery and enrichment values measured by qPCR after every round of mRNA display. **c**) Dataset convergence at the peptide level as gauged by normalized Shannon entropy (S.I. 2.1 for definition). In addition to library 5S5 selection and antiselection experiments, two affinity-based mRNA display selections^49^ are displayed for comparison. **d**) Dataset convergence at the amino acid level as measured by log_2_Y* scores. Amino acid aa in position pos is enriched in the selection dataset compared to the antiselection one if log_2_Y*_aa, pos_ > 0. See also the definitions in the figure header and S.I. 2.1; **c**_aa, pos_ is the number of NGS reads with amino acid aa in position pos in a dataset. **e**) Population-wide analysis of the datasets after 6 rounds of selection/antiselection. Plotted are histograms showing peptide distributions in the S-space. The naïve library is mRNA prior to the first round of selection/antiselection. **f**) CNN classifier accuracy as a function of number of mRNA display rounds. The models were trained on 4.75·10^5^ samples from the respective datasets, using 0.25·10^5^ unseen samples for validation. Multiple rounds of mRNA display lead to cleaner datasets, and hence, more accurate models. **g**) CNN classifier accuracy as a function of training dataset size. The models were trained on round 6 data. Model accuracy scales with the size of the training dataset. **h**) Comparison of the deep CNN classifier against traditional machine learning approaches. Both accuracy and area under receiver operating characteristic (AUROC) metrics are displayed. The use of CNN classifiers leads to better models. AB: AdaBoost; 1NN: nearest neighbor classifier; LR: logistic regression; NN: k-nearest neighbors classifier; RF: random forest classifier; GB: gradient boosting classifier. All classifiers except CNN used scikit-learn (0.24.1) implementations.^67^ **i**) Validation of model predictions against experimental data. 65 validation peptides (bVP1–65; all encoded in library 5S5; see also Table S4) were expressed by the FIT system and treated with LazBF/GluRS/tRNA^Glu^ for 2 h. Reaction outcomes were analyzed by LC-MS as described in S.I. 2.8. Model predictions showed good agreement with the experiment.

Next, we tested whether these substrates could be differentiated in an mRNA display format. Peptide-mRNA/DNA chimeras prepared following the standard techniques (Supporting Information (**S.I**.) 2.2 and 2.3)^49^ were modified under the aforementioned conditions, and, following SAv pulldown, the amount of DNA in either bound or unbound fraction was quantified by qPCR. A large difference in DNA recovery between bAP1 and bAP3 was observed (r; defined as the ratio of DNA in the bound over the unbound fractions; r_bAP1_ = 4.6 vs. r_bAP3_ = 0.008; Fig. S1f), but only when intermediate HA-affinity purification and acetone precipitation steps (aimed to eliminate unreacted biotin-PEG_2_-SH and mRNAs that failed to display peptides) were included (data not shown). Enrichment, defined as the ratio of DNA recovery in the experiment over the negative control (an analogous assay where LazB is omitted from the enzyme mix), also pointed to the enzyme-dependent DNA recovery in the bound fraction (enrichment_AP1_ = 223 vs. enrichment_AP3_ = 1.2; Fig. S1f). Combined, these data indicate that the developed mRNA display pipeline can discriminate LazBF substrates based on their fitness.

With these protocols, we performed six rounds of selection and antiselection for library 5S5 following the established conditions, except starting with round 4 of the selection experiment, the LazBF incubation time was shortened to 15 min to adjust selection pressure. The enrichment values increased between rounds during the selection experiment (Fig. 2a), suggesting that the substrate populations of progressively higher fitness were obtained. In contrast, for antiselection, after the initial decrease in round 2, DNA recovery and enrichment remained relatively constant (Fig. 2b). Next generation sequencing (**NGS**) of the resulting DNA libraries revealed that, even after 6 rounds of selection/antiselection, the substrate populations remained highly diverse and had no convergence at the peptide level, which stands in contrast to traditional affinity-based mRNA display selection workflows (Fig. 2c). To analyze convergence at the amino acid level, we computed Y^*^_aa, pos_ scores as a measure of enrichment for amino acid *aa* in position *pos* in the selection dataset compared to the antiselection one (Fig. 2d). Thus, amino acid *aa* in position *pos* appears to be favored by the enzyme if its log transformed Y^*^ score, log_2_Y^*^, is greater than zero, and disfavored when log_2_Y^*^ < 0. This analysis recapitulated our previous^47,53^ findings: for example, Trp in position 7, i. e., position +1 to the fixed Ser residue, had the highest Y^*^ score (log_2_Y^*^ = 2.53), whereas Asp and Glu, which are known to be disfavored by LazBF,^47,53^ had negative log_2_Y^*^ in every position. Overall, the amino acids around the designed modification site (positions 5 and 7) were subject to a stronger discrimination than those further away from Ser6 (compare position-wise variances of log_2_Y^*^ scores; Fig. 2d). For any library member, a statistical fitness score, **S**, can be computed as the sum of log_2_Y^*^ for every amino acid in the variable region. We found that representing peptides in the S-space is an effective way to perform dataset-wide analysis of substrate populations. For example, consistent with the qPCR results, this analysis revealed (Fig. 2e, S2) that the selection generated a highly enriched substrate subpopulation (1.7σ higher than the naïve library), whereas the antiselection did not, because the antiselection peptides resembled the naïve library members (Δ0.5σ). Altogether, these data suggest that the assay produced enriched yet highly diverse substrate populations suitable for further analysis.

### Development and validation of deep learning models

Next, we turned to the development of a deep learning workflow (Fig. 1c). We sought a scalable and generalizable pipeline to build interpretable models which can facilitate downstream enzymatic studies. After considerable experimentation, we opted for a straightforward data preprocessing routine: NGS data was in silico translated, denoised and demultiplexed, after which the resulting peptide datasets were labelled (S.I. 2.5; Table S3). All selection and antiselection peptides received a label of “1” and “0”, respectively. A number of more sophisticated workflows, which included data preclustering, outlier detection or quantification of relative fitness from NGS read counts, were rejected as they consistently led to models of a poorer performance. Peptide sequences were represented as matrices of positionally encoded amino acid-wise extended connectivity fingerprints (**ECFP**s; S.I. 2.5, Fig. S3 and S4),^59^ a technique that has been recently applied to train models which take peptide sequences as input data.^60,61^ ECFP representations are built directly from the chemical structures of constituent amino acids, and thus, they bypass the limitation of many popular approaches based on biophysical descriptors,^62,63^ which are typically limited to 20 proteinogenic amino acids. At the same time, ECFP representations are more interpretable than one-hot encoding and related techniques. A deep convolutional neural network (**CNN**; 2.5 million parameters; S.I. 2.5 and Fig. S5) was selected as the model architecture, primarily due to its fast training. However, we note, that neither the choice of the model architecture (also tested were recurrent networks, transformers and fully connected networks) nor the nature of peptide representation were particularly critical from the accuracy perspective.

With these methods, we turned to benchmarking the overall workflow. First, we ascertained whether multiple rounds of mRNA display were important by training CNN models on NGS data for each selection/antiselection round using 4.75·10^5^ and 0.25·10^5^ samples for training and validation, respectively (Fig. 2f). Model accuracy increased from 0.823 for round 1 data to 0.992 for round 6, indicating that multiple rounds of mRNA display can furnish progressively cleaner datasets for deep learning. The amount of training data also proved important. Although reasonable models could be trained on as few as 10^2^ peptides from the round 6 dataset (Fig. 2g), the log-log plot of the accuracy vs. the number of training samples was nearly linear, with model accuracy reaching up to 0.997 when 10^7^ peptides were used for training. Notably, saturation in method performance was not observed in either experiment, which suggests that running more rounds of mRNA display and/or increasing the sequencing depth could further improve the accuracy of the method. The latter approach might be particularly straightforward because the throughput of contemporary NGS instruments reaches 10^10^ reads/day.^64^ We also benchmarked our workflow against several traditional machine learning methods (*k*-nearest neighbors, adaptive/gradient boosting, logistic regression and random forest classifiers) and found that deep neural networks were consistently superior (Fig. 2h).

The experiments above evaluated model performance in simple classification tasks where a model is tasked with assigning library 5S5 peptides as belonging to either the selection or antiselection datasets, with NGS data used as the ground truth. In the final experiment, we evaluated whether the models could make more biochemically meaningful predictions, i. e., whether they generalize beyond NGS data and agree with experimentally determined substrate fitness values. To this end, we semi-randomly selected 65 library 5S5 members to ensure a fair test of the model performance (“validation peptide set”, bVP1–65; see S.I. for sequence choices and Table S4) and experimentally investigated their dehydration by LazBF in batch format. The peptides expressed by the FIT system were incubated with LazBF/tRNA^Glu^/GluRS for 2 h under the same conditions as for the mRNA display pipeline. Reaction outcomes were quantified by LC-MS and summarized as modification efficiency values (see S.I. 2.8 for details). The model training pipeline was modified to exclude all validation peptide sequences from the training dataset using Hamming distance ≤ 2 as the cutoff value. Overall, we found that the model predictions tracked the experimental values (Pearson correlation coefficient, ρ_P_ = 0.968; Fig. 2i), indicating that despite being trained as a classifier, the model could also quantify substrate fitness with the mean prediction error of 0.08 ± 0.09 (±σ). The ability to quantify substrate fitness was in line with model’s performance on NGS datasets, that is, the quantification accuracy depended on the amount of training data and the number of mRNA display rounds (Fig. S6a, b). The model excelled at identifying high fitness substrates, whereas underprediction of reaction yields for moderately poor peptides (those with modification efficiencies of ∼0.05–0.15) was the most common source of error.

Altogether, these data demonstrate that the developed mRNA display/deep learning platform can produce accurate models capable of extrapolating substrate fitness across the peptide sequence space. In the subsequent series of experiments, we deployed the model to understand the high-level features of the LazBF substrate space.

### Model-guided population-level analysis of LazBF substrates

In striking contrast to the performance of deep learning models, mRNA display-based statistical metrics such as the S score bore close to no predictive power for the validation peptide set (Fig. 3a). To see whether this is generally true for LazBF substrates, we generated 5·10^6^ random peptides from library 5S5 in silico and estimated their fitness using the model. The analysis of the distribution of model predictions in the S-space demonstrated that statistical enrichments could confidently point to a small fraction of poor substrates (S < –5), but for high fitness peptides the uncertainty of the prediction was too high to be practically useful (Fig. 3b). For example, an average peptide with S = 2.5 had a predicted modification efficiency of 0.49 ± 0.45 (±σ). Representing the outcomes of high throughput enzyme profiling experiments as positional amino acid preferences is a common practice. Based on our results (see also the data for LazDEF below), we argue that at least in some cases such a practice may be counterproductive and misleading, although it remains to be established how general this phenomenon is.

**Figure 3.**
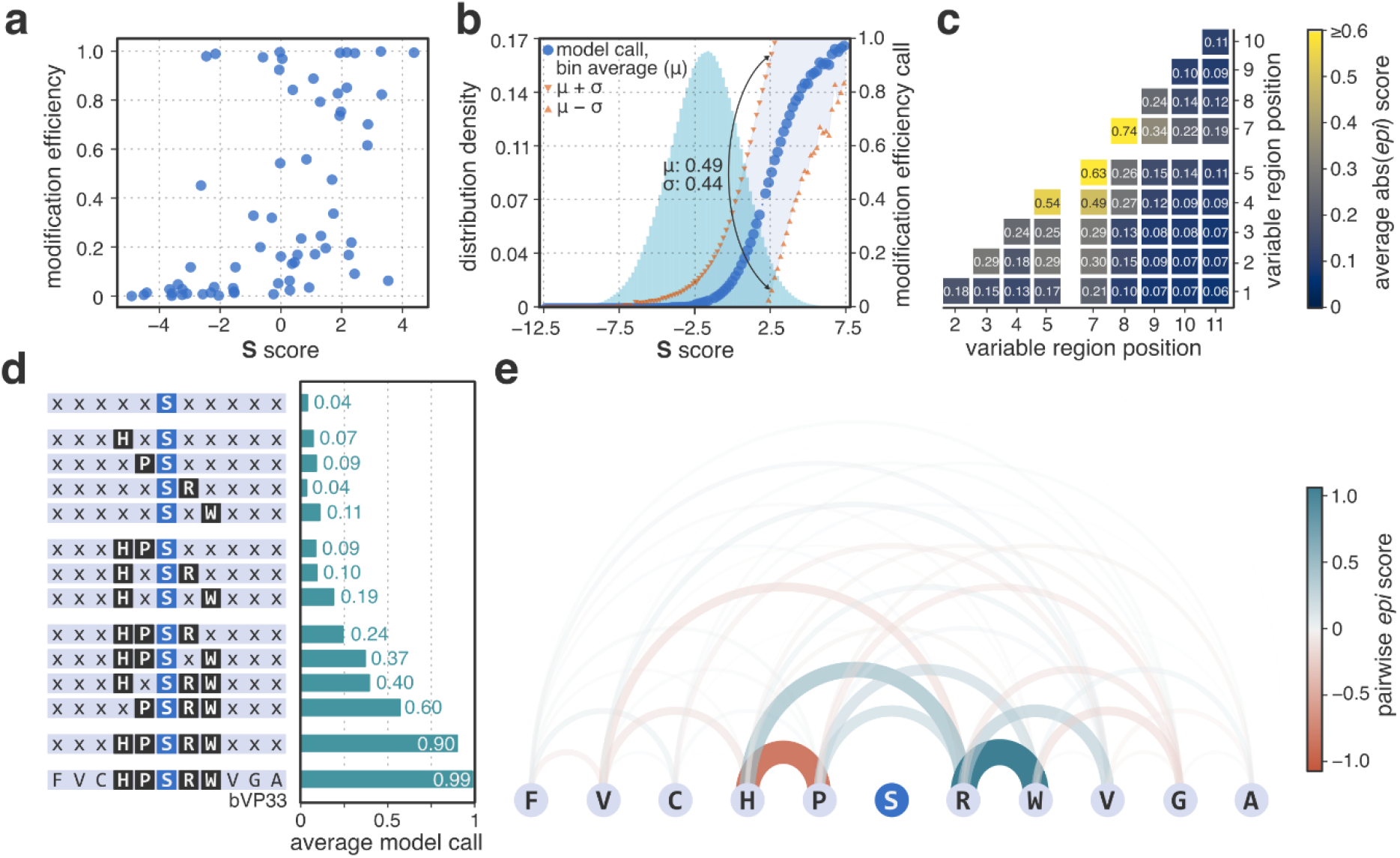
The model enables high-level analysis of LazBF substrate fitness landscapes. **a**) Experimentally measured modification efficiencies of validation peptides (bVP1–65; Table S4) as a function of their S scores. S scores cannot be used to reliably predict fitness of bVP peptides. **b**) Distribution of model predictions in the S-space. Substrate fitness of 5·10^6^ random 5S5 peptides was evaluated with the model. Plotted are binned statistics of model predictions in the S-space. The overall distribution of the peptides in the same space is displayed for reference. The analysis reveals that at best, S scores can be reliably used as anti-determinants of substrate fitness (when S < –5). **c**) Pairwise epistasis between variable positions in the CP of 5S5 peptides. The model was utilized to compute abs(epi) scores using predictions for 5·10^6^ sequences from b). The resulting values can be used to estimate how strongly amino acids in the substrate affect each other’s fitness. Higher values correspond to stronger second order effects. See S.I. 2.1 for computation details. **d**) Analysis of epistatic interactions in bVP33. Average model calls were computed for 2·10^4^ semi-random in silico generated peptides in each case; “x” denotes any amino acid except Ser. **e**) Visualization of all pairwise epistatic interactions in bVP33. Strong epistasis inside the His4-Pro5-Ser6-Arg7-Trp8 motif contributes to the high fitness of the peptide.

Statistically, poor performance of S scores in predicting substrate fitness points to prevalent higher order effects, i. e., the fitness of an amino acid in a given position strongly depends on the rest of the substrate sequence and should not be treated as an independent variable. To quantify some of these effects, we employed the model to compute pairwise epistasis between substrate amino acids in various positions, and summarized the results as *epi* score values (for definition, see S.I. 2.1). A positive *epi* score corresponds to a synergistic effect between amino acids *aa1* and *aa2* in positions *pos1* and *pos2*, respectively. Conversely, a negative *epi* score indicates that on average a substrate containing *aa1* and *aa2* in positions *pos1* and *pos2* has a lower-than-expected fitness if statistical independence of amino acids was assumed (see Fig. S7 for examples). Averaging of absolute *epi* scores over *aa1* and *aa2* can be utilized to estimate how strongly *pos1* and *pos2* influence each other. This analysis showed (Fig. 3c) that amino acids around the modification site (positions 4, 5, 7 and 8) have stronger pairwise epistasis than those distal from Ser6, although a number of long-range interactions were still noticeable (for example, position 1 to position 7; *epi* = 0.21). Overall, such second order effects dominated the substrate fitness landscape for LazBF, which explains why simple amino acid enrichment metrics had limited predictive power. For instance, validation peptide bVP33 underwent near quantitative dehydration by LazBF (0.97) due to the presence of His4-Pro5-Ser6-Arg7-Trp8 motif. Multiple pairwise epistatic interactions within the motif (Fig. 3d and 3e) facilitated substrate fitness despite the low statistical fitness score (S = 0.04), and no single amino acid was solely responsible for the high modification efficiency.

The diversity and abundance of epistatic interactions in LazBF substrates suggest that the enzyme likely makes extensive but transient contacts with the substrate’s CP during the two-step catalysis. Despite the variety of high fitness substrates, LazBF is less promiscuous than it might appear, as only 4% of library 5S5 peptides were predicted to undergo efficient dehydration (Fig. 3d). In a number of preliminary experiments, we investigated whether LazB preferences could be explained by the propensity of the substrates to form specific secondary structures, but the efforts so far have been inconclusive.

### Model-guided peptide-level analysis of LazBF substrates

Integrated gradients (**IG**s) are a popular method for interpreting predictions of deep learning models.^65^ As an attribution technique, IGs seek to understand how individual input features affect a particular prediction by the model. Because in our case peptides are represented as a matrix of ECFPs, IGs can be projected onto the chemical structures of input sequences to visualize model attributions at an atomic resolution. We found this technique insightful in two ways. First, IG attributions facilitated the assignment of PTM sites. For several validation set peptides containing multiple Ser residues in the CP region, the treatment with LazBF yielded chromatographically homogeneous singly dehydrated species (see bVP17, 25, 37 and 51 in Fig. S8a, S9a, 4a, and S10a, respectively), pointing to selective modification of one Ser residue. For bVP37, the model attributed its high score prediction (0.985) primarily to Ser10 (Fig. 4b), while Ser6 was deemed less important, suggesting that the dehydration occurred at the former residue. Tandem mass-spectrometry (MS/MS) of dehydrated bVP37 unambiguously located the modification site to Ser10 (Fig. 4c), and similar analysis performed for bVP17, 25, and 51 confirmed that IG attributions can be utilized to predict modification sites (Fig. S8, S9, and S10). Second, the technique could also be leveraged to dissect the contributions of individual amino acids toward the overall substrate fitness. For several validation set peptides, amino acid-wise IG attributions designated a single amino acid, often far from the modification site (Fig. 4d, S11), as the major reason for a poor dehydration efficiency. Indeed, single-point mutations at specified amino acids improved the experimentally observed substrate fitness in every case.

**Figure 4.**
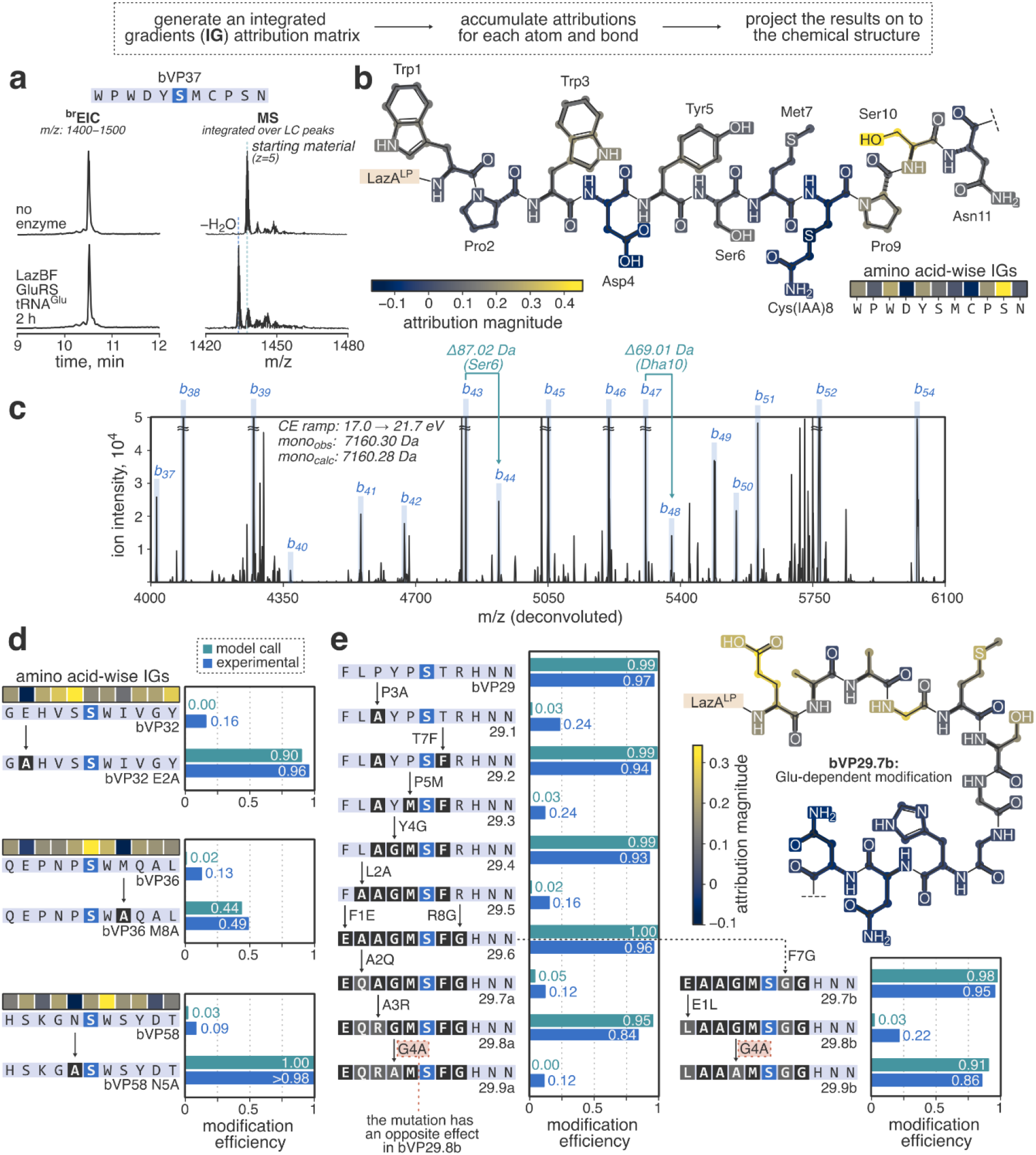
Model-guided dissection of the substrate preferences of LazBF. **a**) LC-MS analysis of bVP37 dehydration by LazBF [a broad extracted ion chromatogram (^br^EIC) and a composite MS spectrum integrated over substrate-derived peaks showing the overall product distribution; see S.I. 2.8 for LC-MS details]. **b**) Atom- and bond-wise accumulated IG attributions for bVP37. The model suggests that Ser10 is the primary determinant of the high modification efficiency. **c**) A zoomed-in section of a charge-deconvoluted CID fragmentation spectrum for singly dehydrated bVP37; *y*-ion assignments and neutral molecule losses are omitted for clarity. The spectrum allows unambiguous assignment of the dehydration site to Ser10, consistent with the model’s suggestion. See Fig. S8–10 for more examples. **d**) Amino acid-wise IGs provide an intuition for relative amino acid contributions to the total substrate fitness. Experimentally measured increase in modification efficiency for three single-point mutants of bVP32, 36, and 58 underscores the model’s ability to identify amino acids critical for LazBF-mediated dehydration. See Fig. S11 for more examples. **e**) Substrate space traversal study for bVP29 (see also the accompanying text). The model was employed to find a sequence of bVP29 mutants which alter substrate fitness at each step. The route identified by the model was validated experimentally. Collectively, this study points to the complex and unintuitive substrate preferences of LazBF.

The model – together with the aforementioned techniques – enabled a detailed evaluation of LazBF’s catalytic promiscuity. Ultimately, we found that the complexity of the substrate landscape, as hinted at by the analysis of epistatic interactions, precludes reasonable simplifications to a set of straightforward rules. To illustrate the intricacy of LazBF substrate preferences, we performed a sequence space traversal study for one validation set peptide, bVP29. Specifically, we utilized the model to find a chain of mutations which drastically alter substrate fitness at each step (Fig. 4e). The model pointed to numerous inconspicuous amino acid replacements distal from the modification site which either abrogated (for example, L2A mutation in bVP29.4) or restored (A3R in bVP29.7a) LazBF-mediated dehydration at Ser6. Altogether, we found that i) the presence of an aromatic amino acid next to the modification site or elsewhere in the CP is not absolutely required for modification (bVP29.7b and bVP29.9b); ii) even though in general negatively charged Glu/Asp in the CP region strongly decrease substrate fitness, some peptides instead rely on the presence of Glu for dehydration (see E1L mutation in bVP29.7b and the corresponding IG attributions); and iii) analogous mutations in homologous substrates (G4A for bVP29.8a and bVP29.8b) can lead to opposite outcomes.

Given the uncovered complexity of LazBF preferences, and hence the infeasibility of manual annotations of substrate fitness for the enzyme, we argue that the models constructed with our platform represent a powerful tool to facilitate the study of promiscuous lanthipeptide and thiopeptide dehydratases. Our results demonstrate that the substrate preferences for LazBF, as obfuscated as they might be, are discernible, and with enough training data, deep learning can furnish models which are both accurate and generalizable.

### LazDEF profiling

In the final series of experiments, we explored how well the developed platform can be expanded to other PTMs. We chose LazDEF, another component of the lactazole biosynthesis pathway, as the model for this study. LazDE is a YcaO family enzyme^66^ which cyclodehydrates Cys and Ser residues in LazA^CP^ during lactazole biosynthesis (Fig. 5a) to yield thiazolines and oxazolines, respectively.^52^ The dehydrogenase domain of LazF further aromatizes these structures to azoles via FMN-dependent dehydrogenation.^52^ As with LazBF, LazDEF is known to process non-native substrates, but the extent of such promiscuity has not been fully elucidated.^47,53^

**Figure 5.**
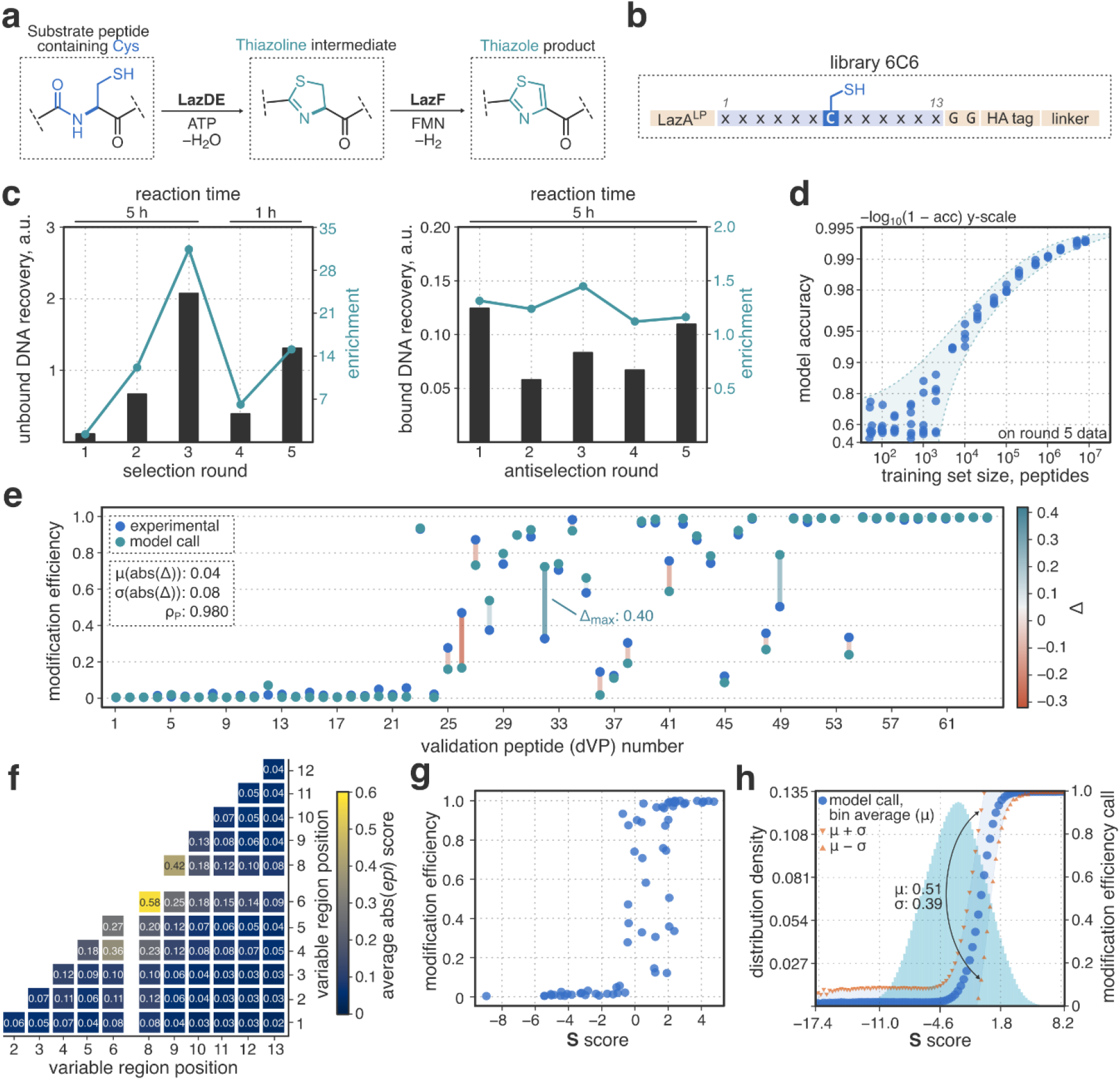
Substrate specificity profiling for LazDEF. **a**) Chemical reactions catalyzed by LazDEF. **b**) Design of the LazDEF substrate library, library 6C6. **c**) Summary of the selection and antiselection experiments. Plotted are respective DNA recovery and enrichment values measured by qPCR after every round of mRNA display. **d**) CNN classifier accuracy as a function of training dataset size. The models were trained on round 5 data. **e**) Validation of model predictions against experimental data. 64 validation peptides (dVP1–64; Table S5) were expressed by the FIT system and treated with LazDEF for 5 h. Reaction outcomes were analyzed by LC-MS as described in S.I. 2.8. Model predictions show good agreement with the experiment. **f**) Pairwise epistasis between variable positions in the CP of 6C6 peptides. The model was utilized to compute abs(*epi*) scores using predictions for 5·10^6^ sequences from panel h). The resulting values can be used to estimate how strongly amino acids in the substrate affect each other’s fitness. Higher values correspond to stronger second order effects. Compared to the results for LazBF, LazDEF substrates are characterized by weaker pairwise epistatic interactions, which aids in explaining the results in panels g) and h). See S.I. 2.1 for computation details. **g**) Experimentally measured modification efficiencies of validation peptides as a function of their S scores. Compared to the LazBF results (Fig. 3a), the S scores for LazDEF substrates prove more informative. **h**) Distribution of model predictions in the S-space. Substrate fitness of 5·10^6^ random 6C6 peptides was evaluated with the model. Plotted are binned statistics of model predictions in the S-space. The overall distribution of the peptides in the same space is displayed for reference. In the interval [–3, 2], which accounts for 46% of the total peptide space, S scores are an unreliable metric of substrate fitness.

To profile the enzyme, we designed mRNA display library 6C6 (Fig. 5b), where the CP region contained a fixed Cys residue flanked by 6 random amino acids on either side. To discriminate LazDEF-modified products (i. e., peptides containing a thiazoline/thiazole residue) from unmodified substrates (peptides bearing Cys6), we opted to use an iodoacetamide-based chemistry to selectively biotinylate the latter (Fig. S12). Thus, the selection protocol was modified to collect and amplify the unbound DNA fraction, while the antiselection amplified SAv pulldown products. In total, we performed five rounds of selection and antiselection (Fig. 5c). Consistent with the LazBF study, the selection recovery and enrichment values increased from round to round except for round 4, when the selection stringency was adjusted, whereas antiselection statistics hovered around the same values. Likewise, the resulting sequence populations had strong enrichments at the amino acid level (Fig. S13), but did not converge at the peptide level (normalized Shannon entropy, H_selection_ = 0.992), which provided an ample amount of training data for deep learning. Training a CNN classifier on round 5 data led to accurate models, where – similar to the LazBF experiments – the model accuracy was proportional to the number of training samples, reaching up to 0.993 for 8·10^6^ input peptides (Fig. 5d, S6c), and deep learning-based classifiers also outperformed traditional machine learning methods (Fig. S14). Validation of model predictions against experimentally measured modification efficiency values for 64 peptides confirmed the ability of the model to generalize beyond NGS datasets (Fig. 5e, Table S5). The LazDEF model predictions were in good agreement with the experimental values [ρ_P_ = 0.980; µ(abs(Δ)) = 0.04 ± 0.08 (±σ)], indicating that the model might be leveraged for quantitative estimation of LazDEF substrate fitness. Taken together, these results show the flexibility of the developed mRNA display platform and its ability to profile PTM enzymes catalyzing diverse chemical reactions.

In contrast to the similar mRNA display outcomes, LazDEF and LazBF had divergent substrate fitness landscapes. The difference mainly manifested in lower pairwise positional epistasis (compare Fig. 5f vs. 3c), and by extension, higher predictive power of statistical fitness scores for LazDEF (Fig. 5g, h). Compared to LazBF, analysis of the substrate preferences for LazDEF through the prism of log_2_Y* values was more meaningful. Consistent with the prior studies,^47,53^ LazDEF primarily relied on amino acids in positions – 1 and +1 to discriminate its substrates, preferring small (Gly/Ser/Ala) amino acids on either side of the modification site, and strongly disfavored Asp/Glu anywhere in the CP (Fig. S13a). However, we note that even in this case, S scores could not reliably predict the fitness of nearly half of the total substrate space: 46% of all library 6C6 substrates had S scores between –3 and 2 where statistical predictions can be inaccurate (Fig. 5h). Accordingly, numerous exceptions to the aforementioned substrate preferences were apparent (Table S5). For example, LazDEF readily modified validation peptide dVP31 (cyclodehydration efficiency: 0.89) which contained Asp adjacent to the modification site (Fig. S15).

## Conclusions

Our study demonstrates that mRNA display can produce large amounts of clean, labelled data for supervised deep learning applications. We found that highly accurate models of enzymatic substrate preferences of two PTM enzymes catalyzing different reactions can be constructed using a unified experimental pipeline. The resulting classifier models could be employed for quantitative assessment of reaction yields, prediction of modification sites, and to analyze the influence of individual amino acids on the overall substrate fitness. The deep learning workflow proved superior to traditional machine learning methods and to statistical enrichment metrics, commonly used for analysis of high throughput enzyme profiling data. Combined, these advances have allowed us to dissect the catalytic preferences of a Ser dehydratase and a YcaO cyclodehydratase, which uncovered unusually complex substrate fitness landscapes in both cases. We believe that the LazBF and LazDEF models will facilitate lactazole bioengineering, and more generally, that the developed platform will foster the study of catalysis by promiscuous PTM enzymes.

## Data availability

Experimental procedures are summarized in Supporting Information and Supplementary Tables. Source Python code for data analysis, trained model weights for LazBF and LazDEF, and instructions to reproduce the work can be found at *https://github.com/avngrdv/mRNA-display-deep-learning*. NGS datasets were deposited to DDBJ (accession number: DRA013287; to be released upon publication). Other data are available from the corresponding authors upon reasonable request.

## Acknowledgements

We thank Yuchen Zhang, Ethan Evans and Adam Beattie for stimulating scientific discussions related to this work. This work was supported by KAKENHI (JP20K15407 to A.V.; JP16H06444 to H.S., Y.G., and H.O.; JP20H05618 to H.S.; and JP20H02866 and JP 19K22243 to Y.G.) from the Japan Society for the Promotion of Science.

