## Supplementary Information for "Accurate models of substrate preferences of post-translational modification enzymes from a combination of mRNA display and deep learning"

### 1. General

Reagents were purchased from Nacalai Tesque, Wako Pure Chemical Industries, Sigma-Aldrich Japan, or TCI Chemicals and were used as received. Anti-HA magnetic beads were from Pierce Thermo Fisher Scientific (#88836). Magnetic beads with immobilized streptavidin (SAv) were from Invitrogen (Dynabeads Streptavidin C1, # 65001). Biotin iodoacetamide probe for Cys labelling was from Sigma Aldrich (CAS: 292843-75-5; #B2059). Biotin–$PEG_2$–SH probe was synthesized as described in section 2.9.

Oligonucleotides for library assembly (HPLC and OPC purification grade for regular and randomized sequences, respectively) were purchased from GeneDesign Inc. (Osaka, Japan). Other primers were from Eurofins Genomics (OPC purification grade). All oligonucleotides were used as received. PCR amplifications were carried out in a BioER TC-96GHBC thermal cycler. qPCR analysis was performed using a LightCycler Nano instrument (Roche) running on LightCycler Nano Software v.1.0.

Protein purification and synthesis of *Streptomyces lactacystinaeus* $tRNA^{Glu}$ were done as previously described.[1–3] Nanodrop 2000c spectrophotometer (Thermo Scientific) equipped with a 5 mm pathlength quartz cuvette was used for protein concentration measurements.

NMR spectra were recorded on a JEOL JNM-ECS400 spectrometer ($^{1}$H NMR at 400 MHz and $^{13}$C NMR at 100 MHz). Dimethyl sulfoxide solvent peak served as the internal standard for $^{1}$H NMR (δ = 2.50) and $^{13}$C NMR (δ = 39.52).

### 2. Methods

#### 2.1. Definitions

**qPCR metrics: DNA recovery and enrichment**. Here, and elsewhere in the text, cDNA refers to “complementary DNA”, which is the first strand product of reverse transcription for an mRNA library. cDNA recovery was defined as

$$\mathbf{r} = \frac{n_{\text{modified}}}{n_{\text{unmodified}}} \qquad (1)$$

where *n* is molar cDNA amount as quantified by qPCR. “Modified” refers to the fraction of the library that was modified by the enzymes (in the case of LazBF, the “modified” fraction was biotinylated, whereas for LazDEF, it was not). Note, that this definition differs from our earlier work.[2] Enrichment of an mRNA display experiment is defined as

$$\text{enrichment} = \frac{\mathbf{r}_{\text{experiment}}}{\mathbf{r}_{\text{negative control}}} \qquad (2)$$

For LazBF, the “negative control” was an assay analogous to the “experiment”, except LazBF was omitted from the reaction mixture during enzymatic treatment. For LazDEF, the “negative control” was an assay with no LazE added to the enzyme premix.

**Normalized Shannon entropy**. Normalized Shannon entropy was calculated to estimate dataset convergence at the peptide level using sequence lists after .fastq parsing and filtration (section 2.5). Here, a “peptide” is the primary sequence of the randomized CP region. Peptide frequency, $\boldsymbol{f}_{\text{pep}}$ in the dataset is

$$\boldsymbol{f}_{\text{pep}} = \frac{\mathbf{c}_{\text{pep}}}{\mathbf{c}_{\text{total}}} \tag{3}$$

where $\mathbf{c}_{\text{pep}}$ is the number of NGS reads in the dataset corresponding to peptide *pep*, and $\mathbf{c}_{\text{total}}$ is the total number of reads in the dataset. Then, normalized Shannon entropy is

$$\mathbf{H}_{\text{dataset}} = -\sum_{\text{pep}} \frac{\boldsymbol{f}_{\text{pep}} \cdot \log_2 \boldsymbol{f}_{\text{pep}}}{\log_2 \mathbf{c}_{\text{total}}} \tag{4}$$

$\mathbf{H}_{\text{dataset}}$ values are bound in the interval [0, 1], where 1 corresponds to a completely random dataset (every entry is unique), and 0 represents peptide lists consisting of a single sequence repeated $\mathbf{c}_{\text{total}}$ times.

**Y* and S scores**. Y* score helps with the dataset analysis at the amino acid level. Analogous to (3), frequency of amino acid *aa* in one of the randomized positions *pos* is:

$$\boldsymbol{f}_{\text{aa, pos}} = \frac{\mathbf{c}_{\text{aa, pos}}}{\mathbf{c}_{\text{total}}} \tag{5}$$

Then, a **Y*** score is defined by analogy to our earlier **Y** score metric[2] used for saturation mutagenesis experiments:

$$\mathbf{Y}^{*}_{\text{aa, pos}} = \frac{\boldsymbol{f}^{\text{ selection}}_{\text{aa, pos}}}{\boldsymbol{f}^{\text{ antiselection}}_{\text{aa, pos}}} \tag{6}$$

For a more uniform perception of the data, we usually log transformed $\mathbf{Y}^{*}_{\text{aa, pos}}$. An **S** score for a peptide *pep* is defined as the sum of $\log_2 \mathbf{Y}^{*}_{\text{aa, pos}}$ scores for every amino acid in the random region:

$$\mathbf{S}_{\text{pep}} = \sum_{\text{aa, pos}} \log_2 \mathbf{Y}^{*}_{\text{aa, pos}} \tag{7}$$

***epi* score**. To analyze the magnitude of synergy (or anti-synergy) between amino acids *aa1* and *aa2* in *pos1* and *pos2* of substrate’s CP, we computed *epi* scores. Technically, *epi* scores can be estimated directly from NGS data. For example:

$$\mathrm{epi}_{\mathrm{aa1,\ pos1,\ aa2,\ pos2}} = \log_2 \frac{\boldsymbol{f}^{\mathrm{selection}}_{\mathrm{aa1,\ pos1,\ aa2,\ pos2}}}{\boldsymbol{f}^{\mathrm{selection}}_{\mathrm{aa1,\ pos1}} \cdot \boldsymbol{f}^{\mathrm{selection}}_{\mathrm{aa2,\ pos2}}} \quad (8)$$

where $\boldsymbol{f}^{\mathrm{selection}}_{\mathrm{aa1,\ pos1,\ aa2,\ pos2}}$ is the abundance (frequency) of peptides containing both *aa1* in *pos1* and *aa2* in *pos2* in the dataset. Positive epi scores represent a synergistic relationship between (*aa1, pos1*) and (*aa2, pos2*): the combination of the two in the selection dataset is observed more frequently than expected if variables were statistically independent. Conversely, negative *epi* values statistically point to a degree of anti-synergy between (*aa1, pos1*) and (*aa2, pos2*). Note, that *epi* scores do not directly report on the fitness of peptides containing (*aa1, pos1*) and (*aa2, pos2*); a highly positive $\mathrm{epi}_{\mathrm{aa1,\ pos1,\ aa2,\ pos2}}$ is not necessarily an indication that the peptides containing (*aa1, pos1*) and (*aa2, pos2*) are excellent substrates.

In practice, many $\boldsymbol{f}^{\mathrm{selection}}_{\mathrm{aa1,\ pos1,\ aa2,\ pos2}}$ values had high sampling errors due to low read numbers $\mathbf{c}_{\mathrm{aa1,\ pos1,\ aa2,\ pos2}}$. More importantly, direct computation from NGS data ignores the heterogeneity of modification sites for many selection peptides, which can strongly affect the results. Therefore, instead of a direct approach, we opted to utilize the models for an indirect computation of *epi* scores.

To this end, we generated $5 \cdot 10^6$ library peptide sequences in silico (library 5S5 for LazBF and library 6C6 for LazDEF) containing a single modifiable residue each. For example, for LazBF the peptides were of the form xxxxxSxxxxx, where x is any amino acid other than Ser. Model-derived substrate fitness values for these peptides can be used to compute $\boldsymbol{f}^{\mathrm{selection}}_{\mathrm{aa,\ pos}}$ and hence *epi* as follows:

$$\boldsymbol{f}^{\mathrm{selection}}_{\mathrm{aa,\ pos}} \equiv \mathbf{p}(\mathrm{aa,\ pos} \mid \mathrm{G}) \quad (9)$$

$$\mathrm{epi}_{\mathrm{aa1,\ pos1,\ aa2,\ pos2}} = \log_2 \frac{\mathbf{p}(\mathrm{aa1,\ pos1} \cap \mathrm{aa2,\ pos2} \mid \mathrm{G})}{\mathbf{p}(\mathrm{aa1,\ pos1} \mid \mathrm{G}) \cdot \mathbf{p}(\mathrm{aa2,\ pos2} \mid \mathrm{G})} \quad (10)$$

$$\mathbf{p}(\mathrm{aa,\ pos} \mid \mathrm{G}) = \frac{\mathbf{p}(\mathrm{G} \mid \mathrm{aa,\ pos}) \cdot \mathbf{p}(\mathrm{aa,\ pos})}{\mathbf{p}(\mathrm{G})} \quad (11)$$

$$\mathbf{p}(\mathrm{aa1,\ pos1} \cap \mathrm{aa2,\ pos2} \mid \mathrm{G}) = \frac{\mathbf{p}(\mathrm{G} \mid \mathrm{aa1,\ pos1} \cap \mathrm{aa2,\ pos2}) \cdot \mathbf{p}(\mathrm{aa1,\ pos1} \cap \mathrm{aa2,\ pos2})}{\mathbf{p}(\mathrm{G})} \quad (12)$$

$$\mathrm{epi}_{\mathrm{aa1,\ pos1,\ aa2,\ pos2}} = \log_2 \frac{\mathbf{p}(\mathrm{G} \mid \mathrm{aa1,\ pos1} \cap \mathrm{aa2,\ pos2}) \cdot \mathbf{p}(\mathrm{aa1,\ pos1} \cap \mathrm{aa2,\ pos2}) \cdot \mathbf{p}(\mathrm{G})}{\mathbf{p}(\mathrm{G} \mid \mathrm{aa1,\ pos1}) \cdot \mathbf{p}(\mathrm{G} \mid \mathrm{aa2,\ pos2}) \cdot \mathbf{p}(\mathrm{aa1,\ pos1}) \cdot \mathbf{p}(\mathrm{aa2,\ pos2})} \quad (13)$$

Here, $\mathbf{p}(\mathrm{aa,\ pos} \mid \mathrm{G})$ is the conditional probability that a random peptide bears amino acid *aa* in position *pos* given that it belongs to the selection dataset (G); $\mathbf{p}(\mathrm{aa,\ pos})$ is the probability that a random peptide contains amino acid *aa* in position *pos* and so on. All of the terms in (13) can be directly computed from the modelling experiments above, which provides a uniform estimation of pairwise epistatic interactions between amino acids in the substrate. Several examples of $epi_{aa1,\ pos1,\ aa2,\ pos2}$ score matrices are provided in Fig. S7.

To estimate how strongly *pos1* and *pos2* influence each other, $epi_{aa1,\ pos1,\ aa2,\ pos2}$ values can be averaged over *aa1* and *aa2* as reported in Fig. 3c and 5h:

$$\overline{\left|epi_{pos1,\ pos2}\right|} = \frac{1}{n^2}\sum_{aa1=1}^{n}\sum_{aa2=1}^{n}\left|epi_{aa1,\ pos1,\ aa2,\ pos2}\right| \tag{14}$$

Note, that these values report on the magnitude of the effect, but not its direction (synergistic vs. anti-synergistic), because averaging is done over *absolute* $epi_{aa1,\ pos1,\ aa2,\ pos2}$ values. $\overline{\left|epi_{pos1,\ pos2}\right|}$ can be used to reason about the magnitude of second order effects in the amino acid enrichment study.

**Modification efficiency**. Experimentally measured substrate fitness is reported as modification efficiency calculated as follows:

$$y_{pep} = \frac{\sum A_{modified\ forms}}{\sum A_{total}} \tag{15}$$

Here, A is area under an LC-MS peak; $\sum A_{total}$ is the total area corresponding to substrate-derived peaks, and $\sum A_{modified\ forms}$ is the area under the peaks for PTM-containing peptides. For LazDEF, modified forms usually consisted of thiazoline and thiazole-containing peaks. For LazBF, mixtures of single, double and triple dehydration products were observed in several instances, all of which were integrated to give $\sum A_{modified\ forms}$ values.

#### 2.2. mRNA library construction

**PCR assembly**. DNA libraries were de novo assembled by PCR from oligonucleotide primers (assembly schemes and primer sequences are summarized in Table S1 and S2, respectively). First, overlapping forward and reverse primers were annealed and extended in the primer extension step. To this end, 50 µL of Platinum SuperFi DNA Polymerase solution [PCR buffer (1x), SuperFi Pol (1x), 250 µM each dNTP; **SF PCR mix**; buffer and enzyme from Thermo Fisher] containing 270 nM forward and 250 nM reverse primers was denatured at 95 °C for 60 s. Then, three cycles of annealing (50 °C for library 5S5; 53.1 °C for library 6C6; 30 s each) and extension (72 °C for 30 s) were carried out. In the next step, the extension product (50 µL) was added to SF PCR mix (950 µL) containing 500 nM forward and reverse primers, and five cycles of amplification ensued (three stage thermal cycling including a denaturing step at 95°C for 20 s, annealing at 57.2°C for 20 s, and extension at 72°C for 30 s). After, the mixture was further incubated at 72°C for 60 s, and cooled to 4°C. The outcomes were evaluated by TapeStation (Agilent Technologies). DNA was first extracted by phenol/chloroform/isoamyl alcohol (25:24:1, saturated with 10 mM Tris (pH 8.0), 1 mM EDTA), and then by chloroform/isoamyl alcohol (24:1).

**Transcription**. Extracted DNA was precipitated with ethanol, washed with 70% ethanol in water (v/v), dissolved in water, and added to the transcription reaction mix [40 mM Tris buffer (pH 8.0) supplemented with 20 mM $MgCl_2$, 10 mM DTT, 1 mM spermidine, 0.01% Triton X-100, 240 nM T7 RNA polymerase, 0.04 U/µL RNasin RNase inhibitor (Promega), and 3.75 mM each NTP; 1 ml total volume]. Transcription was allowed to proceed at 37 °C for 14 h, after which 30 µL of 1 unit/µL RQ1 RNase-free DNase (Promega) was added, and the reactions were further incubated for 60 min at 37°C. Reactions were quenched by the addition of EDTA (final concentration [**f.c.**]: 67 mM) and NaCl (f.c.: 270 mM). The transcripts were precipitated with isopropanol, washed with 70% ethanol (v/v), redissolved in water, and purified by 6% polyacrylamide gel electrophoresis (PAGE) containing 6 M urea. RNA extracted from the gel with 300 mM NaCl were collected by ethanol precipitation, dissolved in water, and frozen at –80 °C for storage.

Starting round 2 of selection/antiselection experiments, transcription was scaled down to a 25 µl reaction, and PAGE purification was not performed.

**Puromycin ligation.** mRNA library was attached to puromycin via a Y-ligation performed with the use of T4 ligase. The reaction containing 1 µM mRNA, 1.5 µM puromycin linker and 1 µM T4 ligase in ligation buffer (40 mM Tris, pH 7.8, 10 mM $MgCl_2$, 10 mM DTT, 0.5 mM ATP in a water/DMSO mixture [8:2, v/v]) was incubated at 25 °C for 45 min. Ligated mRNA was extracted with phenol/chloroform as described above, precipitated with ethanol, washed with 70% ethanol, and redissolved in water. Success of puromycin ligation was judged by PAGE (6%, 6 M urea). Ligation product was diluted with water to 6 µM and frozen at –20 °C for storage. This product was used for in vitro translation in the display assay (sections 2.3 and 2.4).

#### 2.3. LazBF profiling

**Translation and reverse transcription**. An *in vitro* translation system was reconstituted by mixing purified ribosome, enzymes, and translation factors.[4,5] The final reaction mixture contained 50 mM HEPES-KOH (pH 7.6), 100 mM KOAc, 2 mM GTP, 2 mM ATP, 1 mM CTP, 1 mM UTP, 20 mM creatine phosphate, 12 mM $Mg(OAc)_2$, 2 mM spermidine, 2 mM DTT, 1.5 mg/mL *E. coli* total tRNA (Roche), 1.2 µM ribosome, 0.6 µM MTF, 2.7 µM prokaryotic IF1, 0.4 µM IF2, 1.5 µM IF3, 10 µM EF-Tu, 10 µM EF-Ts, 0.26 µM EF-G, 0.25 µM RF2, 0.17 µM RF3, 0.5 µM RRF, 4 µ/mL creatine kinase, 3 µg/mL myokinase, 0.1 µM pyrophosphatase, 0.1 µM nucleotide-diphosphatase kinase, 0.1 µM T7 RNA polymerase, 0.73 µM AlaRS, 0.03 µM ArgRS, 0.38 µM AsnRS, 0.13 µM AspRS, 0.02 µM CysRS, 0.06

µM GlnRS, 0.23 µM GluRS, 0.09 µM GlyRS, 0.02 µM HisRS, 0.4 µM IleRS, 0.04 µM LeuRS, 0.11 µM LysRS, 0.03 µM MetRS, 0.68 µM PheRS, 0.16 µM ProRS, 0.04 µM SerRS, 0.09 µM ThrRS, 0.03 µM TrpRS, 0.02 µM TyrRS, 0.02 µM ValRS, 500 µM each proteinogenic amino acid and 1.2 µM puromycin-conjugated library 5S5 mRNA. The mixtures were first incubated at 37°C for 40 min, and then at 25°C for 10 min before adding EDTA (pH 8.0) to a final concentration of 17 mM. Quenched translation reactions were further incubated at 37°C for 10 min to ensure complete dissociation of mRNA from the ribosome. Reverse transcription was performed with MMLV (H–) reverse transcriptase (Promega) using b.t.c.R26 primer at 42 °C for 60 min following manufacturer's protocol (Tris 50 mM, pH 8.3, 2 mM $MgCl_2$).

**Enzymatic treatment and Dha biotinylation**. Cys residues present in the CP of some library 5S5 peptides interfere with the selection scheme due an intramolecular Michael addition between Cys and Dha (non-enzymatic lanthipeptide formation). Such a cyclization precludes proper biotinylation of the dehydration products and biases the outcomes of the assay. To circumvent this issue, prior to initiating the enzymatic reaction, library 5S5 peptides were alkylated with iodoacetamide (**IAA**). Specifically, to one volume of ice-cold reverse transcription product, 0.22 volumes of IAA (10 mg/ml in water) were added on ice. After 2 min, 2.78 volumes of ice-cold LazBF enzyme premix [2.4 µM LazB, 2.4 µM LazF, 1.2 µM *S. lividans* GluRS, and 24 µM *S. lactacystinaeus* $tRNA^{Glu}$ in 60 mM Tris buffer (pH 8.0) supplemented with 12 mM $MgCl_2$ and 6 mM ATP] were added, and the mixture was transferred to a 25°C incubator. The reaction was allowed to proceed for 120 min (selection rounds 1–3, antiselection rounds 1–6) or 15 min (selection rounds 4–6). To stop the reaction, the tube was transferred on ice, because the LazBF activity at 4 °C is negligible.[6] To the resulting product was added Tris (pH 8.5; f.c.: 50 mM) and biotin-$PEG_2$-SH (nominal f.c.: 10 mM, but about half is consumed to alkylate excess IAA from the previous step; added from 200 mM DMSO stock) on ice. The solution was mixed and left on ice for 16–19 h.

**Library purification**. Prior to HA affinity purification, excess thiol was alkylated with IAA (f.c.: 5 mM) for 15 min on ice. Then, one volume of 2x blocking buffer [TBS-T supplemented with 2 mg/mL bovine serum albumin and 2 mg/mL yeast RNA, where TBS-T contained 50 mM Tris, 150 mM NaCl, 0.2% (v/v) tween-20, pH 7.6] was added to one volume of the library mixture. Anti-HA beads (17 µl of 10 mg/mL bead slurry per pmol of library mRNA) were washed twice with TBS-T, once with 1x blocking buffer and added to the sample. Incubation at 4 °C for 60 min ensued, after which the supernatant was discarded and the beads were washed thrice with TBS-T. Bound peptide-mRNA/cDNA chimeras were eluted from the beads with HA peptide (2 mg/mL in TBS-T; sequence: $NH_2$-YPYDVPDYA-$CONH_2$) by incubating the suspension at 37 °C for 15 min, and collecting the supernatant. The elution step was repeated once more, and the supernatants were combined. HA affinity purification was indispensable to remove peptide-unconjugated and puromycin-unligated mRNA/cDNA, chimeras displaying frameshifted peptides, and other translation side-products.

To exclude the possibility of enriching for the sequences containing Cys–S–S–$PEG_2$–biotin disulfides, the library was then reduced with TCEP. The sample was incubated with a solution of TCEP in TBS-T (pH 7.6; f.c.: 5 mM) on ice for 30 min. Finally, to completely remove the traces of biotin-$PEG_2$-SH probe which interferes with the following pulldown, acetone precipitation was carried out. To one volume of the sample, three volumes of ice-cold acetone were added, and the mixture was incubated at –20 °C for 30 min. After, peptide–mRNA/cDNA chimeras were recovered by centrifugation (15300 g for 15 min at 2 °C). The supernatant was discarded, and the pellets were redissolved in TBS-T.

**SAv pulldown**. Biotinylated peptide–mRNA/cDNA chimeras (i.e., the substrates that underwent LazBF-mediated dehydration) were separated from non-biotinylated ones (i.e., those which were not modified) with the SAv pulldown. To one volume of the sample from above, one volume of 2x blocking buffer was added. Streptavidin C1 SAv Dynabeads (17 µl of 10 mg/mL bead slurry per pmol of library mRNA) were washed twice with TBS-T, once with 1x blocking buffer and added to the sample. Incubation at 4 °C for 10 min ensued, after which the supernatant was collected as “unbound” fraction, and the beads were washed twice with TBS-T containing 1 M urea and then once with TBS-T. Elution of the “bound” cDNA was carried out by heating the beads suspended in elution buffer [SuperFi PCR buffer (1x), 250 µM each dNTP, 250 nM T7g10M.F46 and b.t.c.R26 primers] at 95 °C for 5 min.

**PCR amplification**. Concentrations of recovered cDNA, as well as cDNA recovery **r** [eq (1)] were determined by qPCR. Recovered sample aliquots were amplified [10 mM Tris (pH 8.4), 50 mM KCl, 0.1% (v/v) Triton X-100, 2.5 mM $MgCl_2$, 250 µM each dNTP supplemented with 250 nM T7g10M.F46/b.t.c.R26 primers, Taq DNA polymerase (1x) and Sybr Green I (1x; Thermo Fisher)] and the outcomes were analyzed against a six-point calibration curve generated with a naïve library cDNA sample of known concentrations. In addition, analogous analysis of intermediate sample aliquots also enabled to gauge the success of individual steps of the assay.

Thermal cycling to recover library DNA was performed based on the outcomes of qPCR so as to avoid cDNA overamplification. For “bound” fraction, SuperFi Pol (f.c.: 1x) was added to the elution product. For “unbound” fraction, 5 µl of the SAv pulldown supernatant was added to 195 µl of SF PCR mix containing 250 nM T7g10M.F46/b.t.c.R26 primers. In both cases, three stage thermal cycling ensued (denaturing at 95°C for 15 s, annealing at 61°C for 15 s, and extension at 72°C for 15 s), after which DNA isolation, transcription and puromycin ligation steps were performed as described in section 2.2.

**NGS sequencing**. Tailed PCR was used to install Rd1 and Rd2 adapter sequences to library 5' and 3'-ends, respectively. cDNA recovered as described above was PCR-amplified in the SF PCR mix with appropriate primers (primers lists can be found in Table S2 and S3). The product was carried forward to the second PCR step, which used SuperFi Polymerase and Nextera XT v2 Set primers (sequences from Illumina) to install sequencing barcodes on each sample. The success of PCR was evaluated by 3% agarose gel electrophoresis and TapeStation. After, PCR products were combined and column-purified with a NucleoSpin kit (TaKaRa) adhering to manufacturer's protocol. Concentration of the sample was measured with Qubit (Thermo Fisher) using the dsDNA BR kit. cDNA was then appropriately diluted and denatured with 200 mM NaOH per Illumina's protocol. Denatured library [10 pM containing 10–50% (mol/mol) PhiX Control v3 (Illumina)] was sequenced on Illumina's MiSeq instrument in the single read 1x151 cycle mode using v3 chip, collecting data in the .fastq format. Naïve library and selection/antiselection round 2 samples were analyzed on a v2 chip in the 1x301 cycle mode instead. NGS results were deposited to DDBJ (accession number: DRA013287; data to be released upon publication).

#### 2.4. LazDEF profiling

Overall, the assay for LazDEF closely followed the scheme outlined in section 2.3 for LazBF. Assembly of peptide–mRNA/cDNA chimeras was conducted in the same way, except library 6C6 and v.t1.R36 primer were utilized for translation and reverse transcription, respectively. The protocols differed in details of the enzymatic treatment and biotinylation as described below.

**Enzymatic treatment and Cys biotinylation**. To one volume of ice-cold reverse transcription product were added 2.78 volumes of ice-cold LazDEF enzyme premix [2.4 µM LazD, 2.4 µM LazE and 2.4 µM LazF in 60 mM Tris buffer (pH 8.0) supplemented with 12 mM $MgCl_2$, 1.2 mM DTT, and 6 mM ATP], and the mixture was transferred to a 25°C incubator. The reaction was allowed to proceed for 300 min (selection rounds 1–3, antiselection rounds 1–5) or 60 min (selection rounds 4–5), after which the tube was placed on ice. To allow for quantitative conjugation of unmodified Cys residues in library 6C6 substrates, TCEP (f.c.: 2.5 mM, added as a solution in TBS, pH 7.3) was added to the mixture, and the reduction was allowed to proceed for 30 min on ice. Then, after adding biotin-IAA (**BIAA**; nominal f.c.: 10 mM, but about half is consumed to alkylate TCEP from the previous step; from 100 mM DMSO stock), the sample was mixed well, and the conjugation was performed for 120 min on ice.

HA affinity purification, acetone precipitation and SAv pulldown steps were done according to the protocol from section 2.3, except no additional TCEP reduction was performed, and

25 µl (rather than 17 µl) of C1 SAv Dynabeads slurry per pmol of library mRNA was used for pulldown. qPCR analysis and the following cDNA amplification also adhered to the protocol from above, except primers T7g10M.F46 and v.t1.R36 were used for PCR and qPCR.

#### 2.5. NGS data processing and deep learning

**.fastq parser**. Original python code can be found at *https://github.com/avngrdv/mRNA-display-deep-learning* and *https://github.com/avngrdv/FastqProcessor*. Briefly, .fastq data files containing NGS base calls were parsed to retrieve DNA sequences, which were in silico translated. The resulting peptide lists were filtered to discard sequences of incorrect length, ORFs missing stop codons, peptides containing ambiguous symbols, or not conforming to the library design criteria (i.e., those not having fixed sequences flanking the random region on either side). Additionally, the presence of a fixed modifiable residue in the middle of the random CP (Ser6 for LazBF, Cys7 for LazDEF) was asserted, and constant region sequences were trimmed. For LazBF, LazA$^{CP}$ positions 1–11 were kept after trimming; for LazDEF – positions 1–6 and 8–13 (i. e., Cys7 was "excised" and only the variable regions were carried forward). Finally, for each remaining entry, sequencing Q scores corresponding to the variable region were inspected, and the reads containing any Q scores below 30 were discarded.

In some cases, the final datasets were obtained by concatenating peptide lists from different sequencing runs (LazBF selection/antiselection rounds 5 and 6). For naïve library 5S5 and LazBF selection/antiselection round 2 samples, i. e., those sequenced using a v2 reagent kit, Q score thresholds were set to 20 instead of 30. Parser statistics are summarized in Table S3.

**Data preprocessing**. To prepare training and test datasets, a selection and an antiselection peptide list for a particular experiment (for example, LazBF round 6) were merged in the following way. First, the lists were demultiplexed by removing intraset duplicate sequences, and then the entries appearing in both selection and antiselection datasets were discarded altogether. The resulting peptide lists were compared against the validation peptide set (Table S4 and S5) consisting of either 65 (LazBF) or 64 (LazDEF) peptides. Any entry within Hamming distance ≤ 2 (LazBF) or Hamming distance ≤ 3 (LazDEF) from any of the respective validation peptides was discarded. The remaining datasets were labelled (all selection peptides labelled as 1, antiselection as 0), merged, shuffled, and split into train and test sets.

Finally, the peptides were represented as matrices of ECFPs for model training. To this end, an ECFP feature matrix was generated using chemical structures of component amino acids. For LazDEF, 20 proteinogenic structures were utilized; for LazBF, due to the IAA alkylation of Cys residues prior to the enzymatic reaction (section 2.3), Cys was represented as its alkylated form; other 19 amino acid residues had the proteinogenic structure. ECFPs were generated with *max_radius* = 4 in both cases (rdkit 2021.03.3 implementation; http://www.rdkit.org), which resulted in matrices of dimensions (20, 216) for LazBF and (20, 208) for LazDEF. One-hot encoded peptide matrices [dimensions: (11, 20) for LazBF and (12, 20) for LazDEF] were then multiplied by the corresponding feature matrix to produce the final representations [dimensions: (11, 216) for LazBF and (12, 208) for LazDEF].

**Model description and training**. A neural network containing Conv1D layers with batch normalization and of different kernel sizes and dilation rates (12 convolutional layers in total; for model architecture see Fig. SX) was constructed in tensorflow 2.4.1. The final models contained $2.55 \cdot 10^6$ (LazBF) or $2.66 \cdot 10^6$ (LazDEF) parameters and utilized $1.0 \cdot 10^7$ (LazBF) or $8 \cdot 10^6$ (LazDEF) peptides for training. The resulting model weights for LazBF and LazDEF can be found at *https://github.com/avngrdv/mRNA-display-deep-learning*.

The Adam optimizer[7] with parameters $\beta_1 = 0.9$, $\beta_2 = 0.98$ and $\varepsilon = 10^{-9}$ was used for training. The learning rate was varied during the course of training following the Noam schedule with *warmup_steps* = 4000.[8] Dropout[9] (0.25 for LazBF; 0.20 for LazDEF) was applied to regularize the data, and batch size was set to 2048. Binary crossentropy was used as the loss function, and the training was allowed to proceed either for 300 epochs or until validation loss stopped improving (no decrease in 8 epochs). With this setup, the models could be trained in 1–3 days on a single GeForce RTX3090 GPU (Nvidia).

The training of traditional machine learning models (as listed in Fig. 2h) was done with out-of-the-box classifiers implemented in scikit-learn 0.24.1.[10] Peptides were represented either as one-hot encodings or as ECFP matrices, and in general, no large discrepancies between the two representations were observed.

#### 2.6. Design of validation set peptides

Validation set peptides were designed to provide a fair test of model's performance. We argue that the use of completely random peptides is suboptimal from this perspective, as most such sequences are poor substrates (see the data from Fig. 3d – only 4% of the total peptide space is predicted to be high fitness substrates for LazBF), which leads to a biased test. Instead, we opted for a semi-random sampling to select a set that uniformly covers the underlying S-space, which included many “difficult-to-predict” sequences (those where S scores had little predictive power). Briefly, $1 \cdot 10^4$ library 5S5 (or 6C6) peptides were in silico generated. Overly hydrophobic (>5 hydrophobic amino acids) or positively charged (>3 positively charged amino acids) peptides as well as the substrates containing multiple Cys residues were discarded for practical reasons. The remaining peptides were sorted by their S scores, binned and sampled. Overly repetitive sequences were discarded, and bAP1–3/dAP1–3 validation peptides were appended to their respective lists to give the final validation sets, which are summarized in Table S4 and S5.

#### 2.7. Preparation of synthetic DNA for in vitro translation

**PCR**. Linear double-stranded DNA containing a T7 promoter sequence and ORFs encoding library 5S5 or 6C6 substrates were assembled by PCR from synthetic single-stranded DNA oligonucleotides using Taq polymerase. All PCR were performed in 10 mM Tris (pH 8.4), 50 mM KCl, 0.1% (v/v) Triton X-100, 2.5 mM $MgCl_2$, 250 µM each dNTP supplemented with 500 nM of appropriate primers and Taq DNA polymerase. Three stage thermal cycling included a denaturing step at 95°C for 40 s, annealing at 52°C for 40 s, and extension at 72°C for 40 s. The list of all oligonucleotides and assembly schemes can be found in Tables S1 and S2.

In a two-step PCR assembly, template DNA (1:2000 dilution) was first amplified in Taq PCR solution for 9 cycles, followed by a 13-cycle amplification of the PCR product from the first step (1 µL in 200 µL Taq PCR solution). Assembly outcomes were analyzed by 3% agarose gel electrophoresis stained with ethidium bromide. DNA was isolated following phenol/chloroform extraction and ethanol precipitation steps as in section 2.2. Templates prepared in this way were used for in vitro translation without concentration adjustment or further purification.

#### 2.8. Enzymatic reactions in batch format

**In vitro translation and enzymatic reactions**. Template DNA encoding individual library clones were added (20% v/v) to the transcription-coupled in vitro translation premix described in section 2.3. Translation was allowed to proceed at 37 °C for 50 min, after which the mixtures were transferred on ice. To one volume of the translation product, were added five volumes of the enzyme premix (either LazBF/GluRS/$tRNA^{Glu}$ or LazDEF), composition of which was identical to that described in sections 2.3 and 2.4. Reactions were incubated at 25 °C for 120 min (LazBF) or 300 min (LazDEF). After, the tubes were transferred on ice, and one volume of 30 mM iodoacetamide in methanol was added. To precipitate protein and nucleic acid, the mixtures were incubated on ice for 15 min, followed by a 25 °C incubation for another 10 min. The samples were centrifuged (15300 g for 5 min), and 2–8 µL of the supernatant was analyzed by LC-MS.

**LC-MS and MS/MS analysis**. Reaction outcomes were analyzed using Waters Xevo G2-XS QTof instrument equipped with Acquity I-Class UPLC system. HPLC was done on an Acquity UPLC Peptide BEH C4 column [dimensions: 150 x 2.1 mm; pore size: 300Å ; particle size: 1.7 µm] or an analogous C18 column using 0.1% (v/v) formic acid (FA) in water (solvent A) and 0.1% (v/v) FA in acetonitrile (solvent B) as a mobile phase. Analysis was performed at 60°C and 300 µL/min flow rate with one of three methods listed below.

**Method 1**

*LazBF validation peptide set analysis*

| | |
|---|---|
| 0 – 2 min | 5% B |
| 2 – 15 min | 5 – 65% B, linear ramp |
| 15 – 16 min | 95% B |
| 16 – 18 min | 1% B |

**Method 2**

*LazBF miscellaneous experiments*

| | |
|---|---|
| 0 – 1.5 min | 1% B |
| 1.5 – 16 min | 1 – 61% B, linear ramp |
| 16 – 17 min | 95% B |
| 16 – 19 min | 1% B |

**Method 3**

*LazDEF validation peptide set analysis*

| | |
|---|---|
| 0 – 2 min | 1% B |
| 2 – 17 min | 1 – 61% B, linear ramp |
| 17 – 18.5 min | 95% B |
| 18.5 – 22 min | 1% B |

MS analysis was done in a positive polarity/high sensitivity mode with a 0.3 s scan time. Capillary voltage was set to 700 V; ESI source and desolvation temperatures were 120 and 400°C, respectively. Manufacturer-supplied [Glu1]-Fibrinopeptide B or Leu-enkephalin were used as a lockspray standard for continuous mass axis referencing, and the lockspray setup procedure was performed according to the manufacturer's instructions prior to every run.

For tandem mass spectrometry, a data-dependent acquisition method was used. CID fragmentation was triggered in real time if detected ion intensity exceeded $4 \cdot 10^4$ and z = 5. MS/MS spectra were acquired with a 2 s scan time and parameterized collision energy values. In method 1, collision energies were set to ramp from 6–8 to 38–43 eV over the range of acquired *m/z* values (200 to 2000); in method 2, these values were 6–8 to 30–40 eV, and in method 3, 6–8 to 22–28 eV. Samples subject to MS/MS were reanalyzed three times to acquire MS/MS spectra for each method. All acquired spectra were analyzed, but for brevity, reported are only the most informative ones

LC-MS data was analyzed with MassLynx v.4.1. Product distributions were quantified by integrating areas under LazA-derived peaks. To this end, individual compound areas were calculated by summing areas under extracted ion current (**EIC**) chromatogram peaks for complete charged series, i.e.:

$$A_{pep} = \sum_{z} A_z^{EIC}(pep) \qquad (16)$$

and modification efficiencies were estimated using these values as described in section 2.1.

To facilitate data visualization, reported are broad range extracted ion current ($^{br}$EIC) chromatograms generated as previously reported[1] with *m/z* ±100 or ±50 (±500 or ±250 Da) tolerance window. Briefly, despite methanol precipitation, significant interference by the FIT system-derived small molecules in total ion current chromatograms was frequently observed. However, in general, the interfering compounds had a low molecular weight (<1500 Da), and thus, little interference in the *m/z* region above 1000, where most studied peptides were detected (z=5 in most cases), was observed. Generating $^{br}$EIC chromatograms for translated peptides and their reaction products at *z*=5 with *m/z* ±100 (±500 Da) tolerance window enabled visualization of reaction outcomes with minimal interference from the translation components.

MS/MS assignments were done manually. For a given peptide, a series of possible PTM patterns was generated, and for each of these patterns, ladders of *b*- and *y*-ion series were computed. Calculated ion distributions were compared against experimental spectra, and the ion ladder leading to the best match was used to make spectral assignments.

#### 2.9. Synthesis of biotin-PEG$_2$-SH

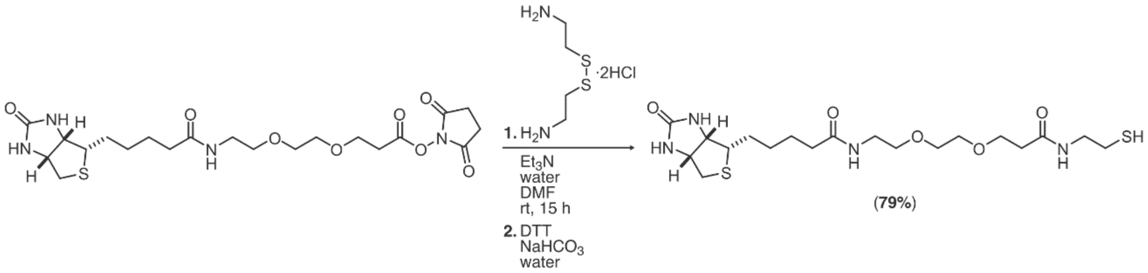

In a round-bottom flask equipped with a magnetic stirring bar, cystamine dihydrochloride (22.5 mg, 0.1 mmol) was dissolved in water (625 µl). To the resulting solution, were sequentially added triethylamine (35 µl, 0.25 mmol), dimethylformamide (1250 µl), and biotin-PEG$_2$-NHS (CAS: 596820-83-6; TCI Chemicals # S0955; 50 mg, 0.1 mmol). The mixture was stirred at room temperature for 15 h, upon which a solution of dithiothreitol (DTT; 150 mg, 0.97 mmol) in saturated aqueous sodium bicarbonate (1500 µl) was added. After stirring for 30 min, another portion of DTT (75 mg, 0.48 mmol) in saturated sodium bicarbonate (750 µl) was added, and the stirring continued for another 30 min. The pH was carefully adjusted to pH 2 with aqueous hydrochloric acid (1 M), and the solution was filtered through a 0.45 µm syringe filter. The product was purified by reverse phase HPLC on a Shimadzu LC-20AP instrument equipped with a Chromolith Prep column (100 x 25 mm, C18 phase; Merck) using 0.1% TFA in water (solvent A') and 0.1% TFA in acetonitrile (solvent B') as a mobile phase at 25 ml/min flow rate. The following gradient was used: 1% B' for 5 min, and then 1 to 36% B' over 35 min. The product eluted at around 22% B' under these conditions. Fractions containing pure biotin-PEG$_2$-SH were combined and lyophilized. The product was obtained as a colorless solid (36.7 mg, 0.079 mmol, 79%).

$^1$H NMR (400 MHz, DMSO-d6) δ 8.01 (t, J = 5.8 Hz, 1H), 7.82 (t, J = 5.6 Hz, 1H), 6.42 (brs, 2H), 4.30 (dd, J = 7.8, 4.1 Hz, 1H), 4.12 (dd, J = 7.8, 4.4 Hz, 1H), 3.59 (t, J = 6.4 Hz, 2H), 3.47 (s, 4H), 3.38 (t, J = 5.9 Hz, 2H), 3.24 – 3.13 (m, 4H), 3.13 – 3.05 (m, 1H), 2.82 (dd, J = 12.5, 5.1 Hz, 1H), 2.57 (d, J = 12.4 Hz, 1H), 2.50 (m, 2H; *overlaps with the solvent peak*), 2.38 – 2.27 (m, 3H), 2.06 (t, J = 7.4 Hz, 2H), 1.68 – 1.55 (m, 1H), 1.55 – 1.39 (m, 3H), 1.38 – 1.19 (m, 2H).

$^{13}$C NMR (101 MHz, DMSO-d6) δ 172.13, 170.17, 162.70, 69.48, 69.45, 69.16, 66.76, 61.03, 59.18, 55.42, 42.06, 39.84, 38.43, 36.10, 35.08, 28.18, 28.03, 25.25, 23.47.

ESI-TOF LC-MS m/z for $[M+H]^+$ ($C_{19}H_{35}N_4O_5S_2^+$); calculated: 463.2043; found: 463.2046.

### 3. Supplementary Figures

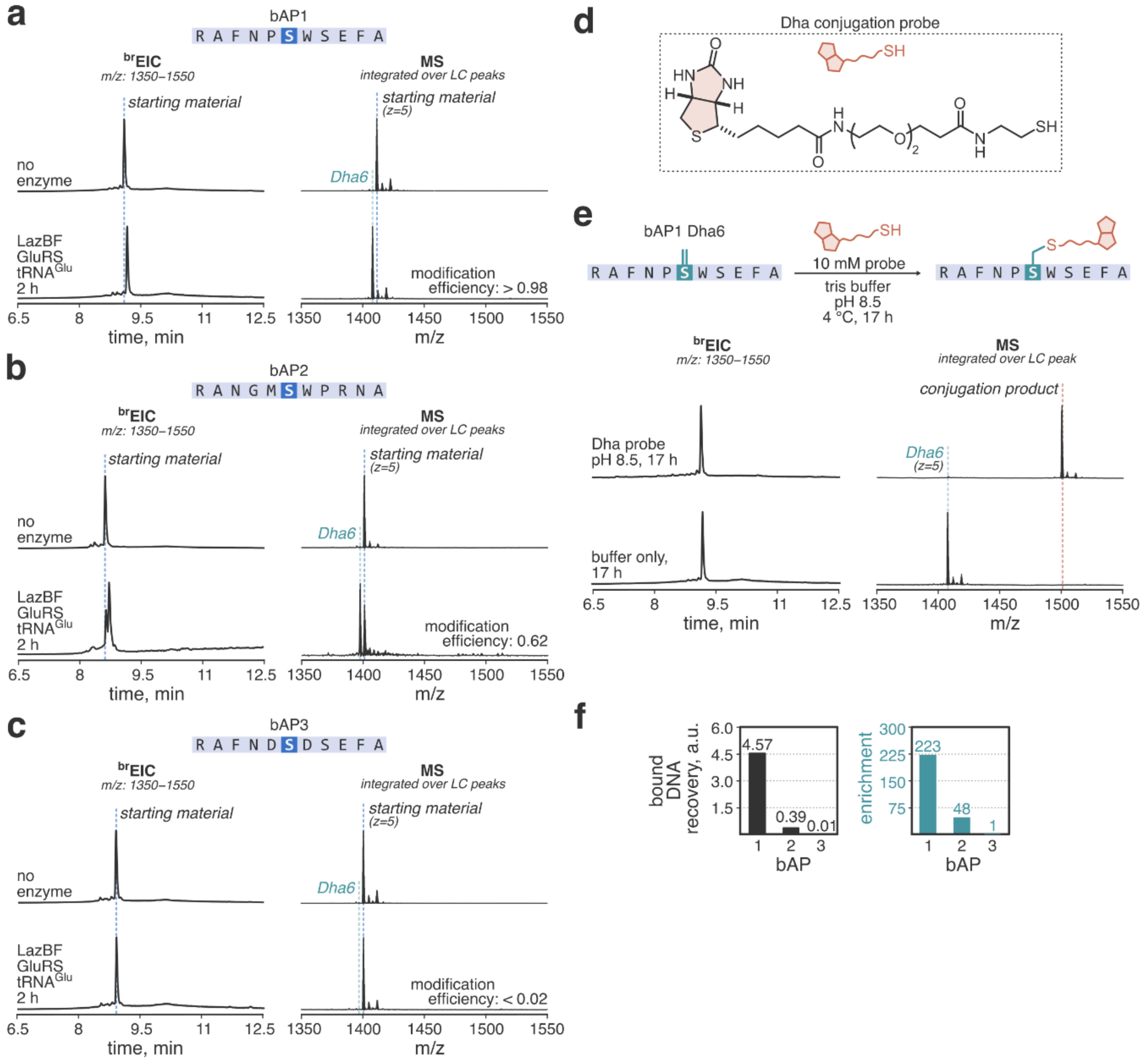

**Figure S1**. Development of the mRNA display assay for LazBF. **a–c**) Modification of bAP1–3 peptides by LazBF as analyzed by LC-MS. Displayed are broad extracted ion chromatograms ($^{br}$EIC) and composite MS spectra integrated over substrate-derived peaks for translation products and LazBF reaction outcomes. See section 2.8 for the details of LC-MS analysis. The three tested peptides (bAP1–3) show differential substrate fitness. **d**) Chemical structure of biotin-PEG$_2$-SH probe for Dha conjugation. **e**) Conjugation of biotin-PEG$_2$-SH to dehydrated bAP1. LazBF reaction product from panel a) was incubated with the probe as specified in section 2.3, and the outcomes were analyzed by LC-MS. Displayed are $^{br}$EIC chromatograms and composite MS spectra integrated over substrate-derived peaks. Clean Dha conjugation takes places under the investigated reaction conditions. **f**) qPCR statistics for the mRNA display assay (section 2.3) performed with bAP1–3. For definition of recovery and enrichment values, see section 2.1. The experiment demonstrates that mRNA display can effectively discriminate the substrates of different fitness.

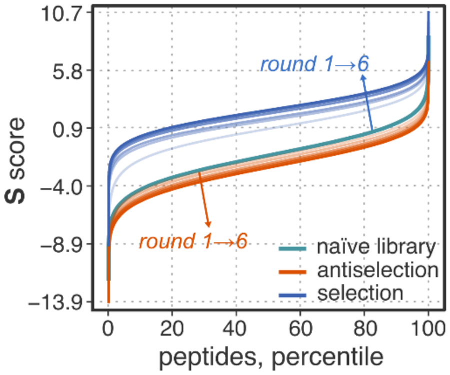

**Figure S2**. Percentile plot showing the distribution of library 5S5 in the S-space during the course of 6 rounds of mRNA display, for both selection (blue lines) and antiselection (orange lines) experiments. The naïve (starting) library is shown in cyan. This analysis confirms that selection, by comparison to antiselection, led to more enriched substrate populations.

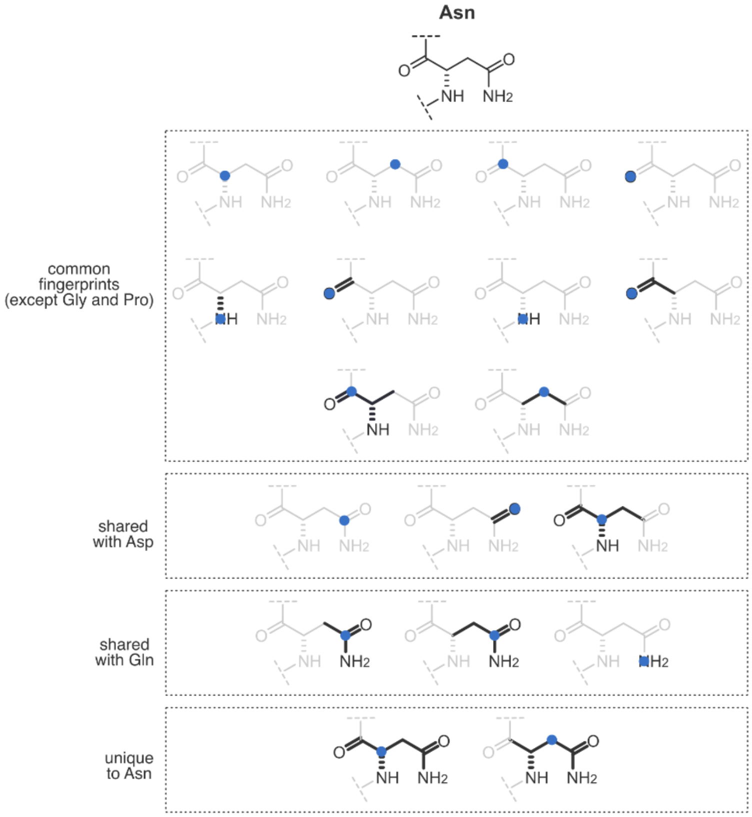

**Figure S3**. Analysis of ECFPs resulting from asparagine at *max_radius* = 4. Blue circles represent the center atom of each fingerprint. ECFPs capture chemical similarity between related amino acids. For example, in this case, Asn shares several fingerprints with Asp and Gln.

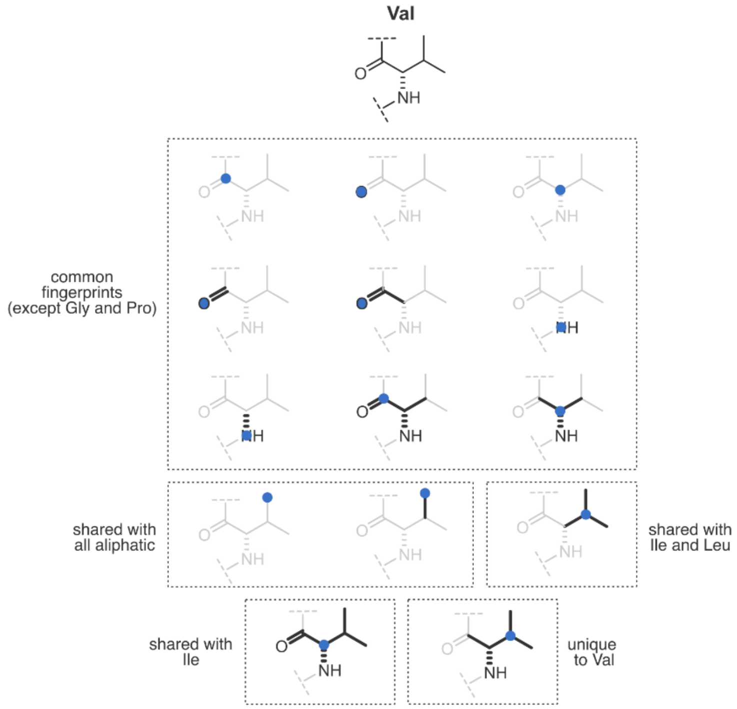

**Figure S4**. Analysis of ECFPs resulting from valine at *max_radius* = 4. Blue circles represent the center atom of each fingerprint. ECFPs capture chemical similarity between related amino acids. For example, in this case, Val shares several fingerprints with other aliphatic amino acids such as Ile and Leu.

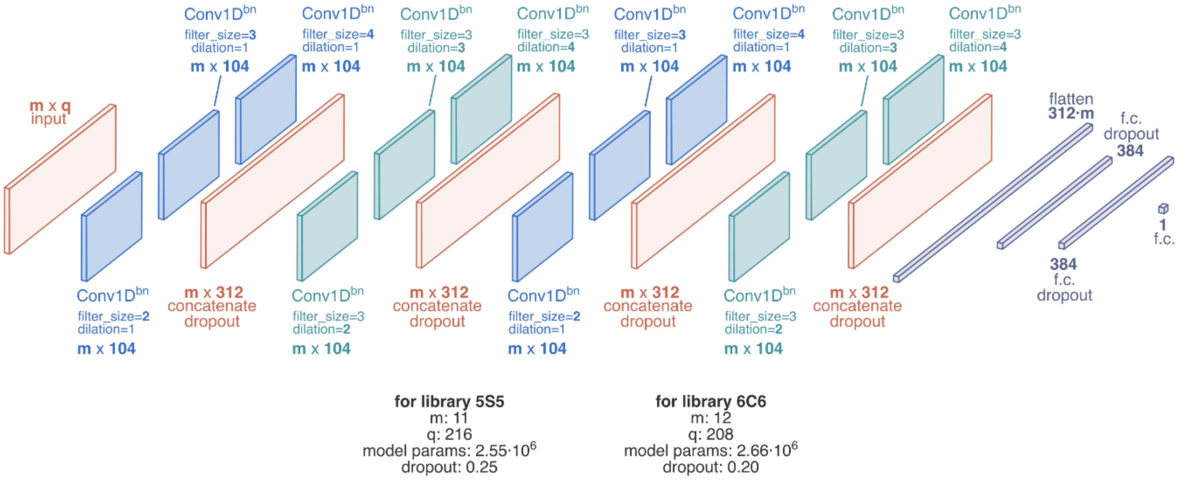

**Figure S5**. Graphical summary of the primary model architecture utilized in this work. Conv1D$^{bn}$: 1D convolutional layer with batch normalization; f.c.: fully connected layer.

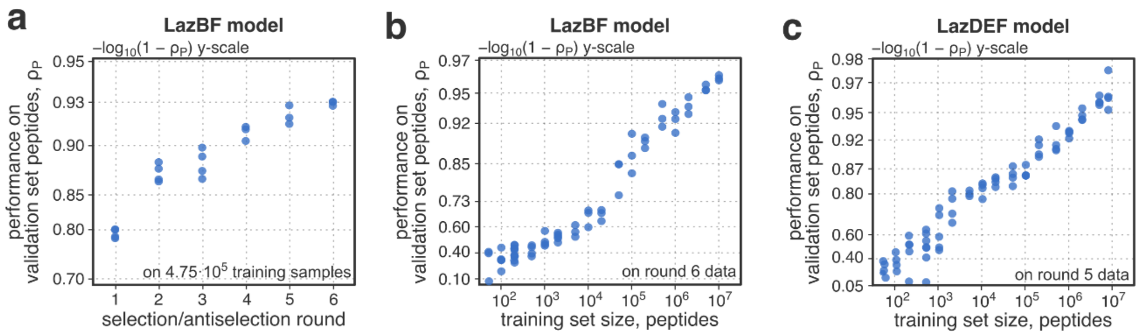

**Figure S6**. Performance of CNN classifiers on validation set peptides. **a**) Pearson correlation coefficient ($\rho_P$) between model predictions and experimentally measured modification efficiencies as a function of number of mRNA display rounds for the LazBF study. The models were trained on $4.75 \cdot 10^5$ samples from the respective datasets. **b,c**) log-log plots of $\rho_P$ vs. training dataset size for LazBF (b; round 6 data) and LazDEF (c, round 5 data) studies. Combined, these data suggest that consistent with the results from Fig. 2f, 2g and 5d, model performance scales with increasing the number of training samples, and that multiple rounds of mRNA display improve accuracy of the resulting models.

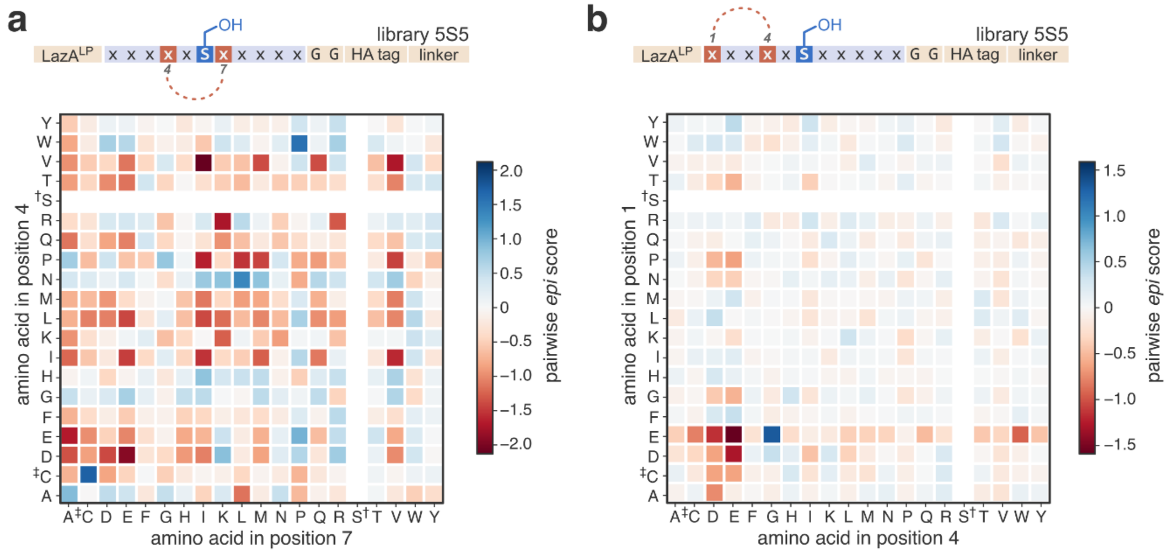

**Figure S7**. Pairwise epistatic interactions in LazBF substrates. See also section 2.1. **a**) Pairwise *epi* scores in library 5S5 peptides between amino acids in position 4 and 7. **b**) Pairwise *epi* scores in library 5S5 peptides between amino acids in position 1 and 4. Note the strong epistasis between Glu1 and Gly4, which was observed and confirmed in the substrate space traversal study (Fig 4e; IG attribution map; peptides bVP29.7b and bVP29.8b). Also compare the magnitude of epistatic effects for positions adjacent vs. distal to the modification site (panel a vs. panel b). †: to avoid the modification site ambiguity, Ser was excluded from the modelling (section 2.1). ‡: Cys was represented as its IAA alkylation product (section 2.5, data preprocessing).

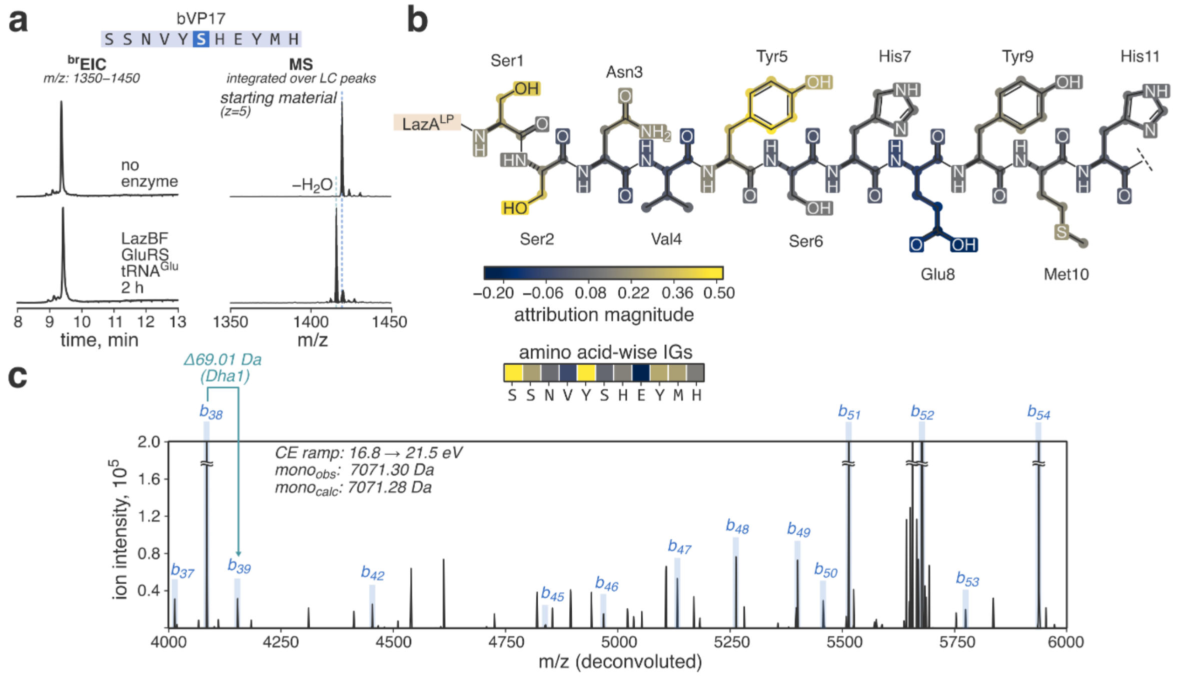

**Figure S8**. IGs help in identifying the sites of LazBF-mediated dehydration in library 5S5 substrates. **a**) LC-MS analysis of the modification of bVP17 by LazBF (a brEIC chromatogram and a composite MS spectrum integrated over substrate-derived peaks showing the overall product distribution; see section 2.8 for LC-MS details). **b**) Atom- and bond-wise accumulated IG attributions for bVP17 as well as amino acid-wise attributions. The model suggests that Ser1 is the primary determinant of high modification efficiency (see amino acid-wise attributions). **c**) A zoomed-in section of a charge-deconvoluted CID fragmentation spectrum for singly dehydrated bVP17; *y*-ion assignments and neutral molecule losses are omitted for clarity. The spectrum allows unambiguous assignment of the dehydration site to Ser1, consistent with the model's suggestion.

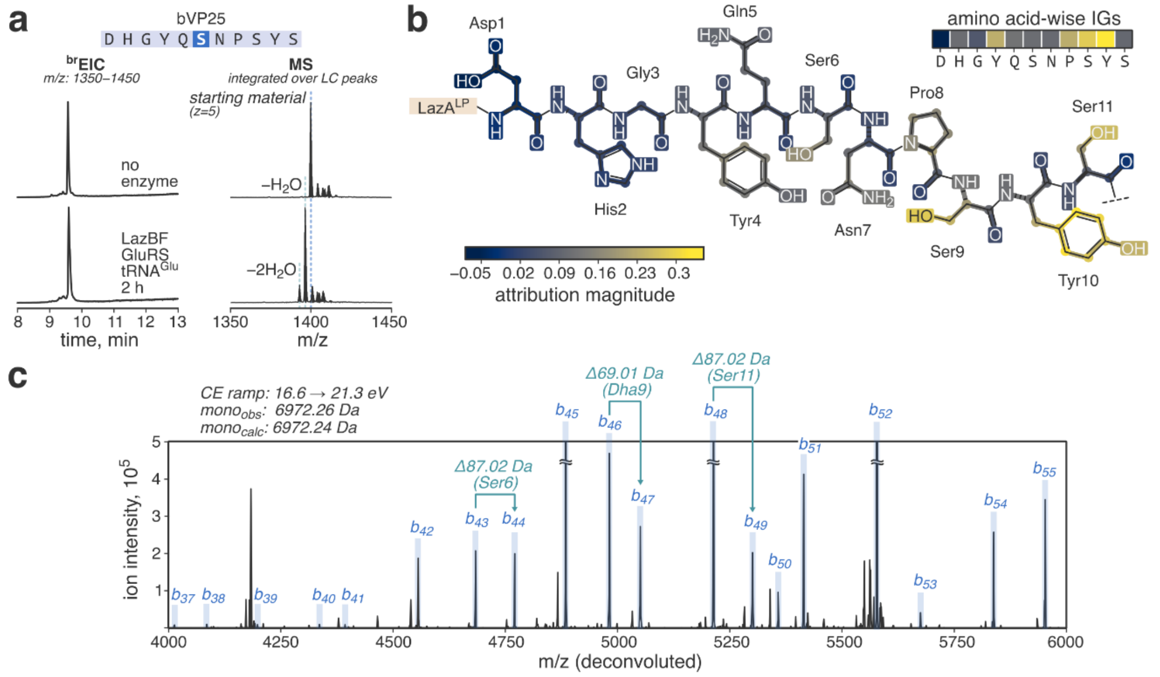

**Figure S9**. IGs help in identifying the sites of LazBF-mediated dehydration in library 5S5 substrates. **a**) LC-MS analysis of the modification of bVP25 by LazBF (a brEIC chromatogram and a composite MS spectrum integrated over substrate-derived peaks showing the overall product distribution; see section 2.8 for LC-MS details). **b**) Atom- and bond-wise accumulated IG attributions for bVP25 as well as amino acid-wise attributions. The model suggests that Ser9 is the primary determinant of high modification efficiency. **c**) A zoomed-in section of a charge-deconvoluted CID fragmentation spectrum for singly dehydrated bVP25; *y*-ion assignments and neutral molecule losses are omitted for clarity. The spectrum allows unambiguous assignment of the dehydration site to Ser9, consistent with the model's suggestion.

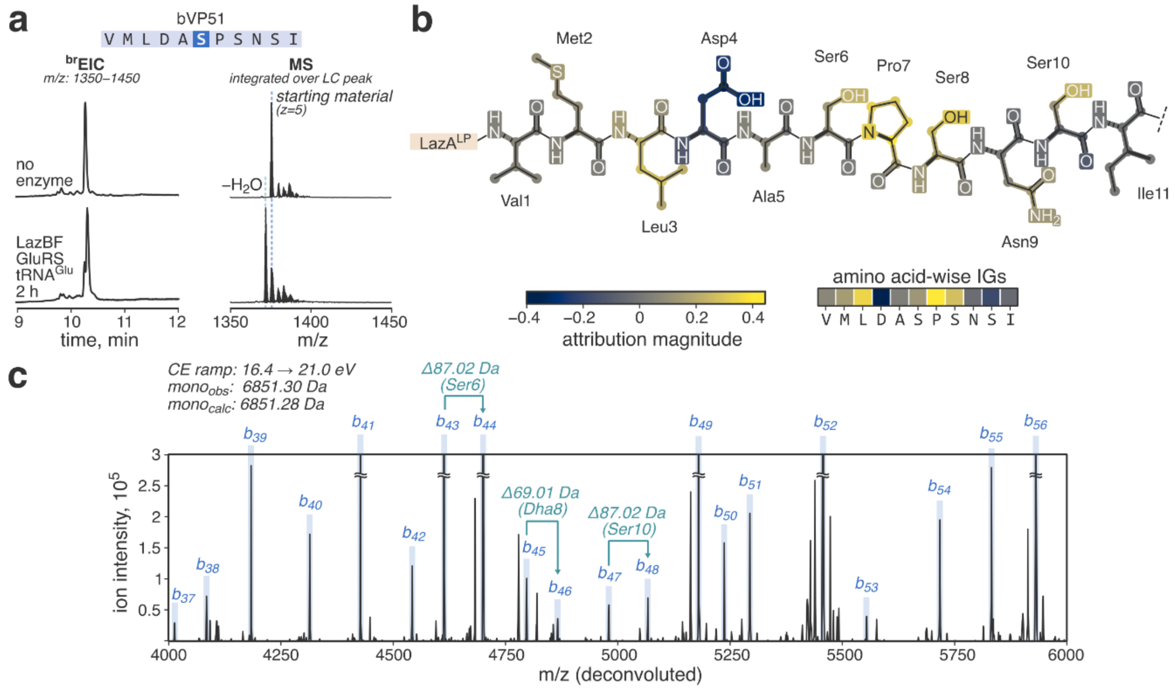

**Figure S10**. IGs help in identifying the sites of LazBF-mediated dehydration in library 5S5 substrates. **a**) LC-MS analysis of the modification of bVP51 by LazBF (a $^{br}$EIC chromatogram and a composite MS spectrum integrated over substrate-derived peaks showing the overall product distribution; see section 2.8 for LC-MS details). **b**) Atom- and bond-wise accumulated IG attributions for bVP51 as well as amino acid-wise attributions. The model suggests that Ser8 is the primary determinant of high modification efficiency. **c**) A zoomed-in section of a charge-deconvoluted CID fragmentation spectrum for singly dehydrated bVP51; *y*-ion assignments and neutral molecule losses are omitted for clarity. The spectrum allows unambiguous assignment of the dehydration site to Ser8, consistent with the model's suggestion.

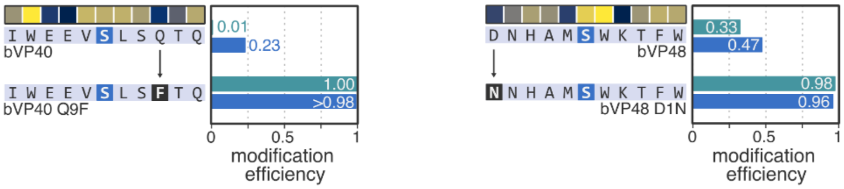

**Figure S11**. Amino acid-wise IGs provide an intuition for relative amino acid contributions to the total substrate fitness. Experimentally measured increase in modification efficiency for two single-point mutants of bVP40 and 48 underscores the model's ability to attribute identify amino acids critical for LazBF-mediated dehydration. See also Fig. 4d.

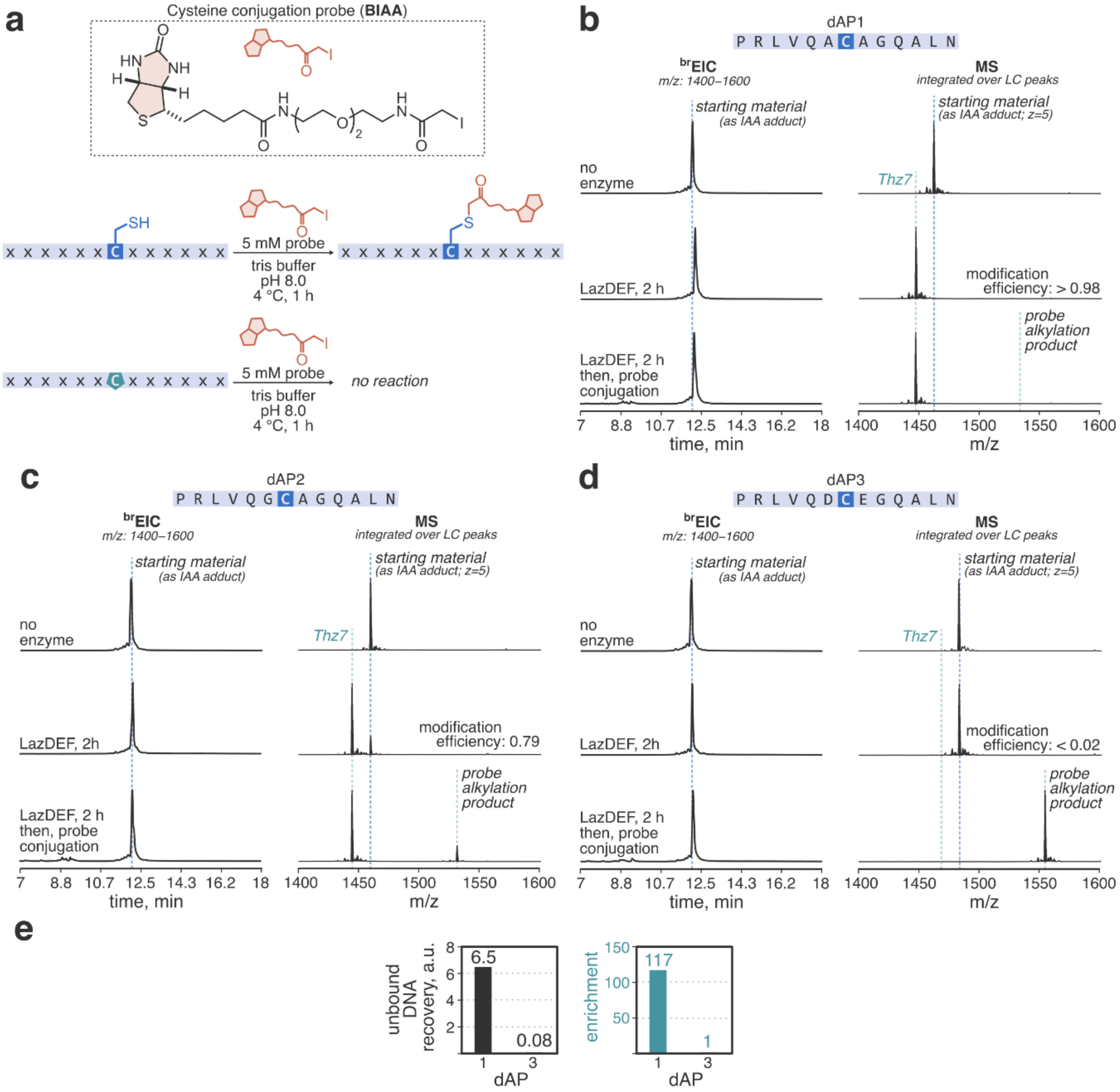

**Figure S12**. Development of the mRNA display assay for LazDEF. **a**) Chemical structure of BIAA probe for Cys conjugation and the labelling scheme. **b–d**) Modification of dAP1–3 peptides by LazDEF and the following BIAA conjugation as analyzed by LC-MS. Displayed are $^{br}$EIC chromatograms and composite MS spectra integrated over substrate-derived peaks after translation, LazDEF reaction, and BIAA conjugation. See section 2.8 for the details of LC-MS analysis. The three tested peptides (dAP1–3) show differential substrate fitness; BIAA selectively can selectively label unreacted peptides. **e**) qPCR statistics for the mRNA display assay (section 2.4) performed with dAP1 and 3. For definition of recovery and enrichment values, see section 2.1. The experiment demonstrates that mRNA display can effectively discriminate the substrates of different fitness.

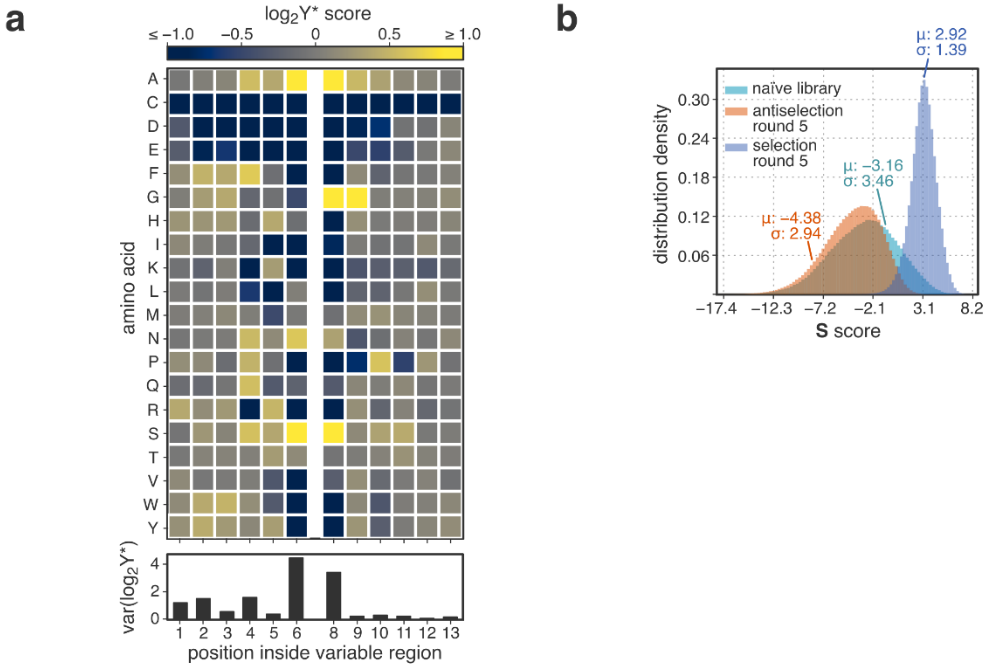

**Figure S13**. Analysis of mRNA display outcomes after 5 rounds of selection/antiselection for the LazDEF study. **a**) Dataset convergence at the amino acid level as measured by $log_2Y^*$ scores. Amino acid *aa* in position *pos* is enriched in the selection dataset compared to the antiselection one if $log_2Y^*$(*aa, pos*) > 0. Also displayed are variances of $log_2Y^*$(*pos*) scores. Compared to the analogous results for LazBF (Fig. 2d), these data indicate LazDEF substrates underwent strong discrimination based on amino acids in positions 6 and 8 (i.e., –1 and +1 relative to the designed modification site). **b**) Population-wide analysis of the datasets after 5 rounds of selection/antiselection. Plotted are histograms showing peptide distributions in the S-space. Naïve library is mRNA prior to the first round of display. Similar to the LazBF study (Fig. 2e), selection experiments produced a peptide population that was more enriched that the one resulting from the antiselection.

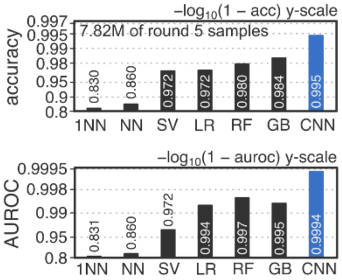

**Figure S14**. Comparison of the deep CNN classifier against traditional machine learning approaches for LazDEF. Both accuracy and area under receiver operating characteristic (AUROC) metrics are displayed. The use of CNN classifiers leads to better models. 1NN: nearest neighbor classifier; NN: *k*-nearest neighbors classifier; SV: linear support vector classifier; LR: logistic regression; RF: random forest classifier; GB: gradient boosting classifier. All classifiers except CNN used scikit-learn (0.24.1) implementations.[10]

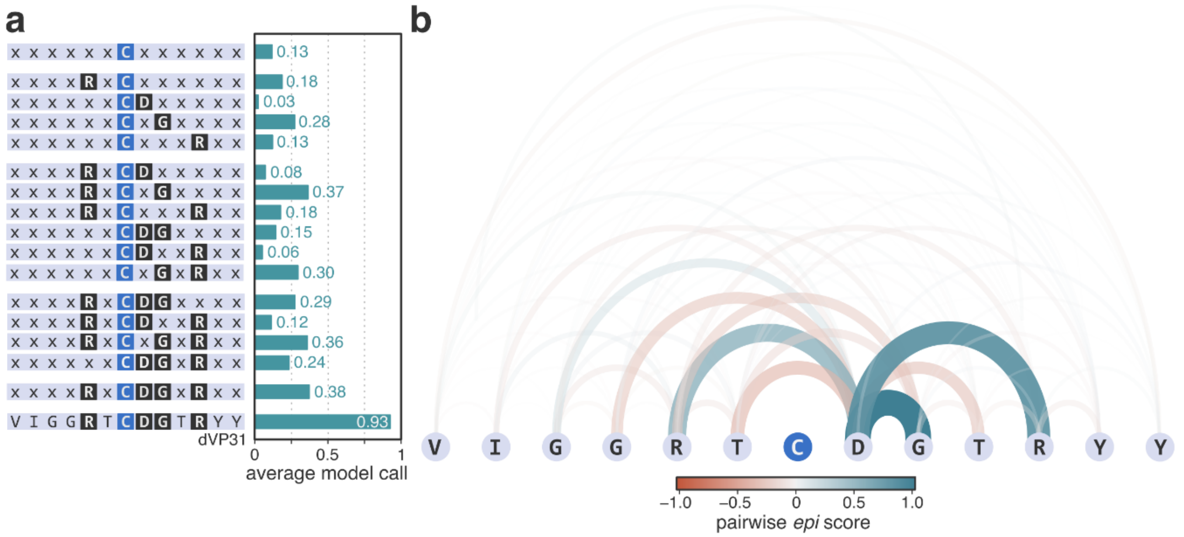

**Figure S15**. Analysis of epistatic interactions in dVP31. d) Average model calls were computed for $2 \cdot 10^4$ semi-random in silico generated 6C6 peptides for each entry; “x” denotes any amino acid except Cys. e) Visualization of all pairwise epistatic interactions in dVP31. Asp8, which usually abrogates LazDEF-mediated modification, has several strong epistatic interactions (Arg5–Asp8, Asp8–Gly9, and Asp8–Arg11) which contribute to high fitness of the peptide.

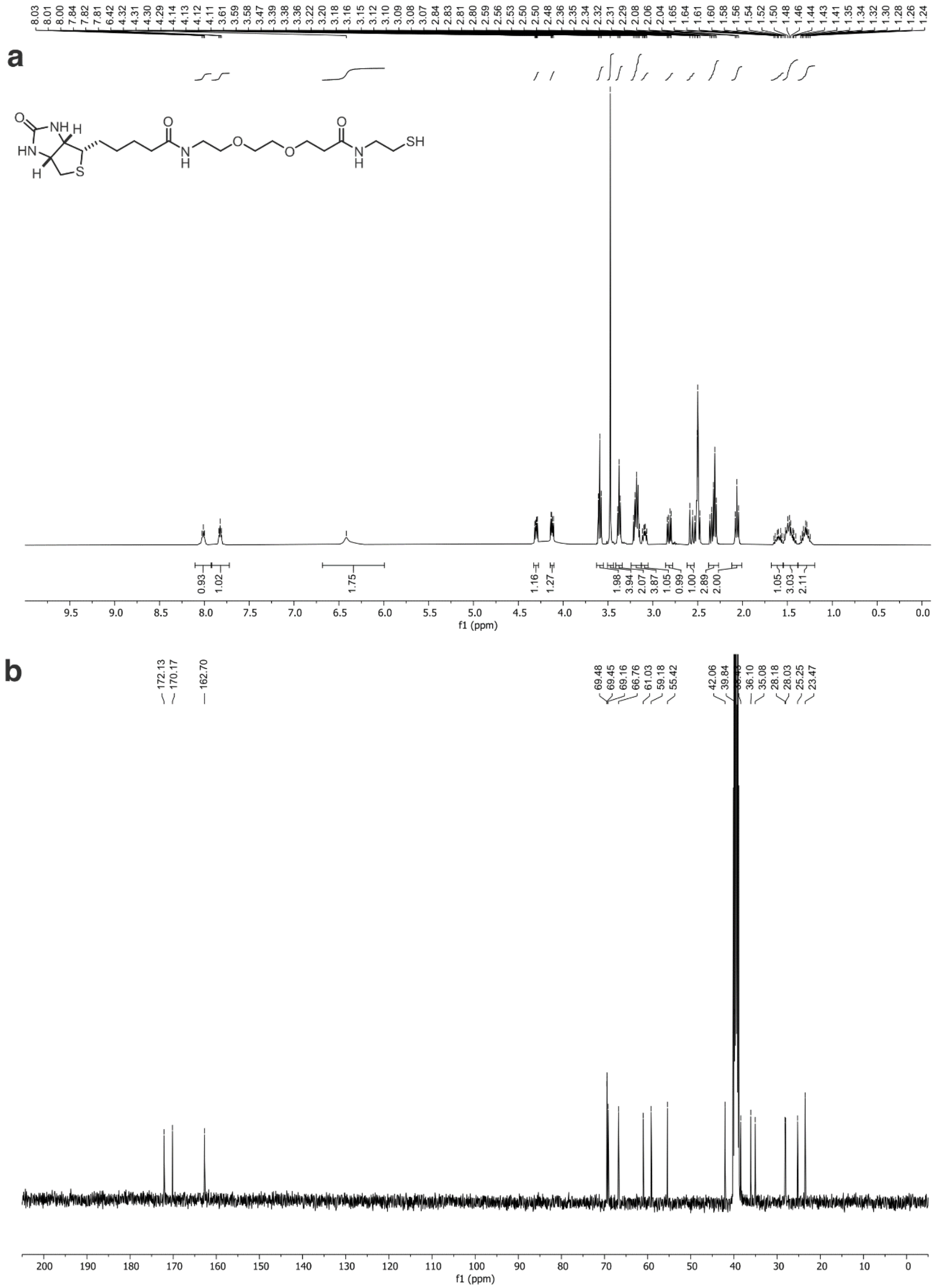

**Figure S16**. NMR spectra [1H (a) and 13C (b)] for biotin–$PEG_2$–SH. See also section 2.9.
